# COPII Sec23 proteins form isoform-specific ER exit sites with differential effects on polarized growth

**DOI:** 10.1101/2020.06.22.165100

**Authors:** Mingqin Chang, Shu-Zon Wu, Samantha E. Ryken, Jacquelyn E. O’Sullivan, Magdalena Bezanilla

## Abstract

COPII, a coat of proteins that form vesicles on the ER, mediates vesicle traffic from the ER to the Golgi. In contrast to metazoans that have few genes encoding each COPII component, plants have expanded these gene families leading to the hypothesis that plant COPII has functionally diversified. Here, we analyzed the gene families encoding for the Sec23/24 heterodimer in the moss *Physcomitrium (Physcomitrella) patens*. In *P. patens, Sec23* and *Sec24* gene families are each comprised of seven genes. Silencing the Sec23/24 genes revealed isoform specific contributions to polarized growth, with the closely related *Sec23D/E* and *Sec24C/D* essential for protonemal development. Focusing on the *Sec23* gene family, we discovered that loss of Sec23D alters ER morphology, increases ER stress, inhibits trafficking to the Golgi and to the plasma membrane in tip growing protonemata. In contrast, the remaining five *Sec23* genes are dispensable for tip growth. While *Sec23A/B/C/F/G* do not quantitatively affect ER to Golgi trafficking in protonemata, they do contribute to secretion to the plasma membrane. Of the three highly expressed Sec23 isoforms in protonemata, Sec23G forms ER exit sites that are larger than Sec23B and Sec23D and do not overlap with Sec23D. Furthermore, ER exit sites labeled by Sec23B or Sec23G form in the absence of Sec23D. These data suggest that *Sec23D/E* form unique ER exit sites contributing to secretion that is essential for tip growing protonemata.

## Introduction

Trafficking from the ER to the Golgi is mediated by the Coat Protein complex II (COPII). COPII facilitates the formation of transport vesicles at ER exit sites promoting the delivery of cargo, including transmembrane proteins, soluble proteins and lipids, to *cis*-Golgi compartments (Brandizzi and Barlowe, 2013; Barlowe and Helenius, 2016; Aridor, 2018; Hanna et al., 2018; Robinson et al., 2015). The COPII coat comprises two layers of protein complexes. The inner layer consists of a Sar1-Sec23-Sec24 lattice, while the outer layer is made up of Sec31-Sec13 heterotetramers (Stagg et al., 2008; Whittle and Schwartz, 2010; Bi et al., 2002; Fath et al., 2007). Sar1 is a small GTPase of the Ras super family that is the master regulator of COPII vesicle biogenesis (Nakano and Muramatsu, 1989). Assembly of COPII coat is initiated by the conversion of Sar1-GDP to Sar1-GTP, which is mediated by Sec12, a guanine nucleotide exchange factor (GEF) that is an ER resident membrane protein (Barlowe and Schekman, 1993; Nakano and Muramatsu, 1989). At the ER surface, activated Sar1 recruits the Sec23-Sec24 heterodimer, which associates with the ER membrane as an “inner coat”. The Sar1-Sec23-Sec24-cargo “pre-budding” complex in turn recruits the Sec13-Sec31 heterotetramer, referred to as an “outer coat” to the nascent bud (Aridor et al., 1998; Salama et al., 1993). The polymerization of Sec13-Sec31 heterotetramers induces membrane curvature and completes vesicle biogenesis (Fath et al., 2007; Stagg et al., 2006; Whittle and Schwartz, 2010).

Selective recruitment of cargos into COPII coated vesicles is driven by ER export signals located on cytoplasmically exposed regions of cargo proteins (Barlowe, 2003). ER export signals of most transmembrane cargo proteins are thought to interact directly with the inner coat components of COPII. However, some transmembrane and most soluble cargo proteins interact with COPII components through ER-resident transmembrane adaptors or receptors (Barlowe, 2003). Sec24 is the COPII component that has been implicated in recognizing these various sorting signals (Mossessova et al., 2003; Nufer, 2003). However, several studies suggested that Sec23 and Sar1 may also play roles in cargo sorting by directly binding to the export signals (Aridor et al., 2001; Mancias and Goldberg, 2007; Giraudo and Maccioni, 2003).

Based on the high sequence similarity among COPII paralogs, and cross-species complementation studies (De Craene et al., 2014; Khoriaty et al., 2018), it has been assumed that functions of COPII components are conserved across eukaryotes. While components of the COPII coat are well studied in budding yeast and mammalian cells, illustrating their molecular functions in plants has been more challenging. Comparative genomic analyses revealed that gene families encoding COPII components have undergone gene expansions in plants with *Sec23* exhibiting the largest expansion (Aridor et al., 2001; Mancias and Goldberg, 2007; Giraudo and Maccioni, 2003; Schlacht and Dacks, 2015). While there are only one and two *Sec23* genes in *S. cerevisiae* and mammals (eg. mice and humans) (Jensen and Schekman, 2011), respectively, there are seven in both *P. patens* and *A. thaliana* (Brandizzi, 2018). Since Sec23 and its heterodimeric partner Sec24 are implicated in cargo sorting, the larger *Sec23* gene family could suggest that plants have more diverse sorting signals requiring specificity at the level of the Sec23/Sec24 heterodimer. Consistent with this, in contrast to yeast, which has a single gene for most COPII components, all other components of COPII in plants are encoded by multiple genes, suggesting that cargo specificity may translate to formation of distinct COPII complexes and vesicle populations. Alternatively, larger gene families could simply indicate that plants have higher functional redundancy in COPII encoded gene families.

Genetic studies in *A. thaliana* have demonstrated that loss of specific COPII components caused developmental defects. A complete knockout of AtSec24A was lethal (Faso et al., 2009; Nakano et al., 2009). However, a missense point mutation in AtSec24A (R693K) resulted in ER morphology defects and disruption of ER-Golgi integrity. Interestingly, these phenotypes could not be rescued by either AtSec24B or AtSec24C (Faso et al., 2009; Nakano et al., 2009), suggesting that AtSec24B and C are functionally distinct from AtSec24A. Even though AtSec24B and AtSec24C did not rescue the *Atsec24a* missense mutant and each were not important for plant viability, they were shown to influence the development of reproductive cells (Tanaka et al., 2013). Functional diversification was also suggested by analysis of AtSec16 mutants in which storage protein precursors were accumulated abnormally in *Atsec16a* null mutants, but not in *Atsec16b* null mutants (Takagi et al., 2013).

While *P. patens* and *A. thaliana* have similar numbers of genes encoding COPII components, moss has several attributes that make it particularly suited to study COPII function. *P. patens* juvenile tissues are haploid and comprise multiple tissue types that establish the plant body. Protonemal tissue, which emerges from the spore and is also readily propagated vegetatively by tissue homogenization, is composed of tip-growing cells constituting a two-dimensional filamentous network. The predictable patterns of growth and cell division provide an easily tractable system for analysis of mutant phenotypes. As a single-cell-layer, protonemal tissue also enables high-resolution microscopy of endogenously tagged fluorescent proteins in live cells (Rensing et al., 2020). In addition to amenable tissue morphology and haploid lifestyle, *P. patens* is amenable to transient RNA interference for rapid identification of gene function and exhibits high rates of CRISPR Cas9-mediated genome editing (Rensing et al., 2020; Mallett et al., 2019). Here, we utilized these molecular genetic and live-cell imaging tools to systematically characterize the *Sec23* and *Sec24* gene families. Analysis of mutant phenotypes revealed a high degree of functional specificity among Sec23 and Sec24 isoforms. Focusing on the *Sec23* gene family, we discovered that Sec23 isoforms differentially influence ER to Golgi trafficking, and secretion to the plasma membrane as well as the size of presumptive ER exit sites.

## Results

### Sec23 and Sec24 isoforms differentially affect tip growth

*P. patens* has seven *Sec23* and seven *Sec24* genes. Based on sequence similarity, we found that there are four subclasses of *Sec23* and *Sec24* genes and in both cases six of the seven genes exist in closely related pairs (Fig 1A, B). For *Sec23*, with the exception of *Sec23A*, at least one member of each subclass is highly expressed in protonemal cells (Fig 1A). For *Sec24*, we found that one of the seven *Sec24* genes (*Sec24E)* was not expressed in protonemata (Fig1B). Of the remaining six genes, *Sec24D* exhibited the lowest expression while the other five Sec24 genes were expressed at similar levels (Fig 1B). To determine if Sec23 and Sec24 participate in protonemal growth, we used transient RNA interference (RNAi) to silence the *Sec23* and *Sec24* gene families. We used a region of the coding sequence from a member of each subclass (Fig1A, B, solid lines under gene models). These sequence regions share a high degree of sequence similarity to the other gene in each subclass (Fig 1A, B, dashed lines under gene models, Table S1) and would effectively simultaneously silence both genes. We did not target *Sec24E* since we were unable to detect a transcript.

**Figure 1.**
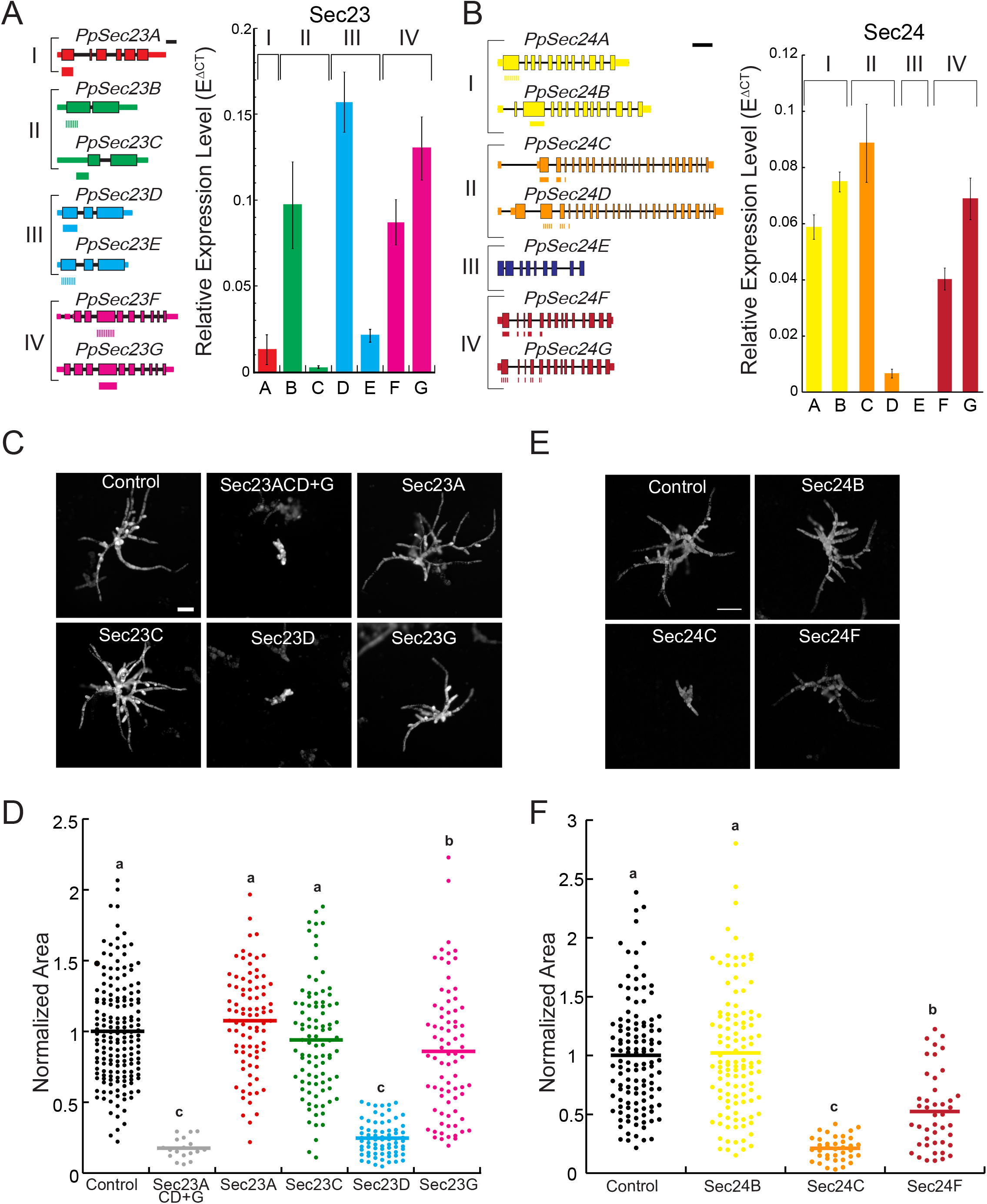
*Sec23* and *Sec24* are required for polarized growth. Gene models of the *P. patens Sec23* (A) and *Sec24* (B) genes are shown with exons indicated by boxes and introns by thin black lines. Coding and untranslated regions are denoted by thick and thin boxes, respectively. The lines underneath the gene models represent sequence regions targeted by RNAi constructs. Scale bar is 500bp. Graphs depict the relative expression of *Sec23* and *Sec24* genes normalized to UBIQUITIN10 in 8-day-old wild-type moss plants regenerated from protoplasts. N = 3, Error bars are s.e.m. (C, E) Representative chlorophyll autofluorescence images of 7-day-old plants regenerated from protoplasts expressing the indicated RNAi constructs. Scale bar, 100 µm. (D, F) Quantification of plant area is based on the area of the chlorophyll autofluorescence and is presented normalized to the area of the control for each experiment. Means are indicated by horizontal lines. Letters indicate groups with significantly different means as determined by a one-way ANOVA with a Tukey *post hoc* test (α=0.05).

By co-transforming the Sec23G RNAi construct together with a construct containing the target sequences for *Sec23A, C* and *D*, we targeted the entire *Sec23* gene family and discovered that silencing the seven *Sec23* genes leads to a dramatic reduction in growth (Fig 1C, D). Surprisingly, we found that silencing subclass III comprised of *Sec23D* and *Sec23E* resulted in a similarly severe growth defect (Fig 1C and D). In contrast, silencing any of the other subclasses did not affect growth (Fig 1C, D). These results demonstrated that there is functional specificity among *Sec23* isoforms, with only subclass III playing a critical role in protonemal growth, while the remaining five *Sec23* isoforms appear to be dispensable.

Similar to silencing subclass III *Sec23* genes, silencing *Sec24C* and *Sec24D* resulted in a 79% reduction in plant area (Fig 1E, E). Silencing *Sec24F* and *Sec24G* also impaired growth, but to a lesser extent resulting in a 48% reduction in plant area (Fig 1E, E). On the other hand, silencing *Sec24A* and *Sec24B* did not affect growth, although both genes were expressed at high levels. These data suggest that like Sec23, there is isoform specificity among *Sec24* genes.

### Sec23 genes in subclass III are functionally redundant

To test whether *Sec23D* and *Sec23E* within subclass III are functionally redundant, we designed RNAi constructs targeting the untranslated regions (UTRs) of each gene (Fig 2A). Targeting the untranslated regions of both *Sec23D* and *Sec23E* resulted in small unpolarized plants with an 85% reduction in plant area as compared to control RNAi plants (Fig 2B, C). Silencing *Sec23D* alone resulted in a 59% reduction in plant area, while silencing *Sec23E* did not affect plant area (Fig 2B, C). Given that *Sec23D* is expressed seven times higher than *Sec23E* (Fig 1B), it is possible that *Sec23D* contributes to protonemal growth more than *Sec23E*. To test whether *Sec23E* can replace *Sec23D* function if expressed at sufficient levels, we performed complementation studies by driving the full-length coding sequence of *Sec23D* or *Sec23E* with the constitutive maize ubiquitin promoter. Interestingly, the reduction in plant area resulting from silencing of *Sec23D* was restored to similar levels when either *Sec23D* or *Sec23E* full-length cDNAs were expressed (Fig 2B, C), indicating that *Sec23D* and *Sec23E* are functionally redundant, and that the highly expressed *Sec23D* is the predominant group III isoform critical for protonemal growth.

**Figure 2.**
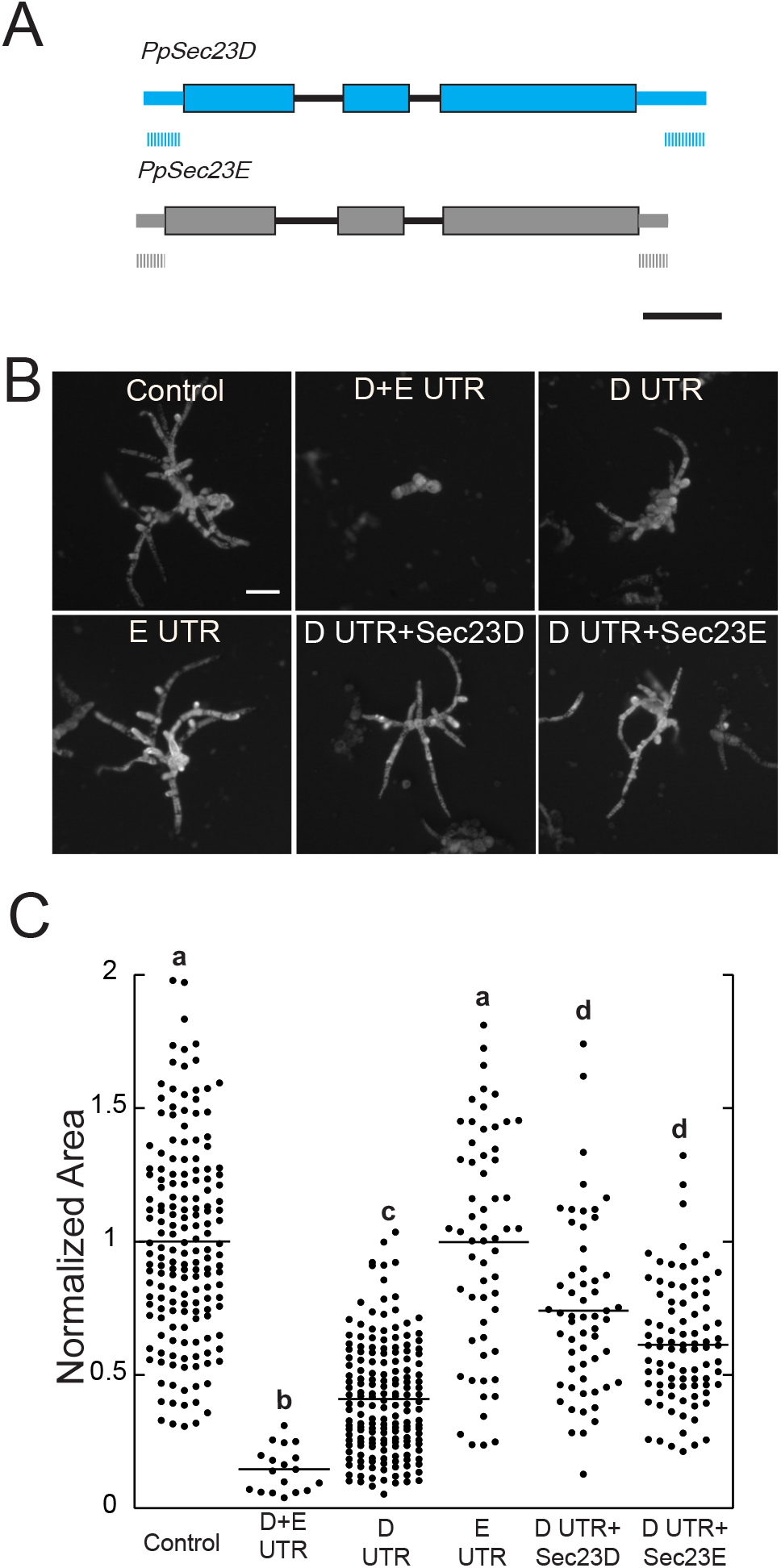
*Sec23D* and *Sec23E* are functionally redundant. (A) *Sec23D* and *Sec23E* gene models are shown with exons indicated by boxes and introns by thin black lines. Coding and untranslated regions are denoted by thick and thin boxes, respectively. The lines underneath the gene models represent sequence regions that were targeted by *Sec23D* or *Sec23E* UTR RNAi constructs. Scale bar is 500bp. (B) Representative chlorophyll autofluorescence images of 7-day-old plants regenerated from protoplasts expressing the indicated constructs. Scale bar, 100 µm. (C) Quantification of plant area was normalized to the control for each experiment. Letters indicate groups with significantly different means as determined by a one-way ANOVA with a Tukey *post hoc* test (α=0.05).

### CRISPR-Cas9-mediated deletion of Sec23 isoforms

Using transient RNAi, we were able to deduce that *Sec23* genes display isoform specific functions during protonemal growth. However transient RNAi assays are limited to phenotypic characterization of 7-day-old plants. To further characterize the molecular functions of *Sec23*, we generated stable loss-of-function mutants of *Sec23* genes using CRISPR-Cas9 mediated genome editing. We targeted each gene with one protospacer located in the first coding exon (Fig S1A, S2). We obtained single and higher-order mutants of *Sec23* genes carrying out-of-frame mutations resulting in premature stop codons in their transcripts (Fig S1A, S2, Table S2). Consistent with the RNAi results, 7-day-old *Δsec23d* plants were 70% smaller than wild type plants while *Δsec23e* plants exhibited no growth defects (Fig 3A, B). We isolated a number of *sec23d* null alleles (Fig S1A, Table S2), which had similar growth phenotypes (Figure S1B). We also found that in three-week-old plants regenerated from protoplasts, *Δsec23d-5* plants were 62.6% smaller than wild type (Fig 3C, D), and Δ*sec23d-5* had fewer gametophores compared to wild type (Fig 3C, E), suggesting that slow growing protonemal tissues resulted in a developmental delay in the production of gametophores.

**Figure 3.**
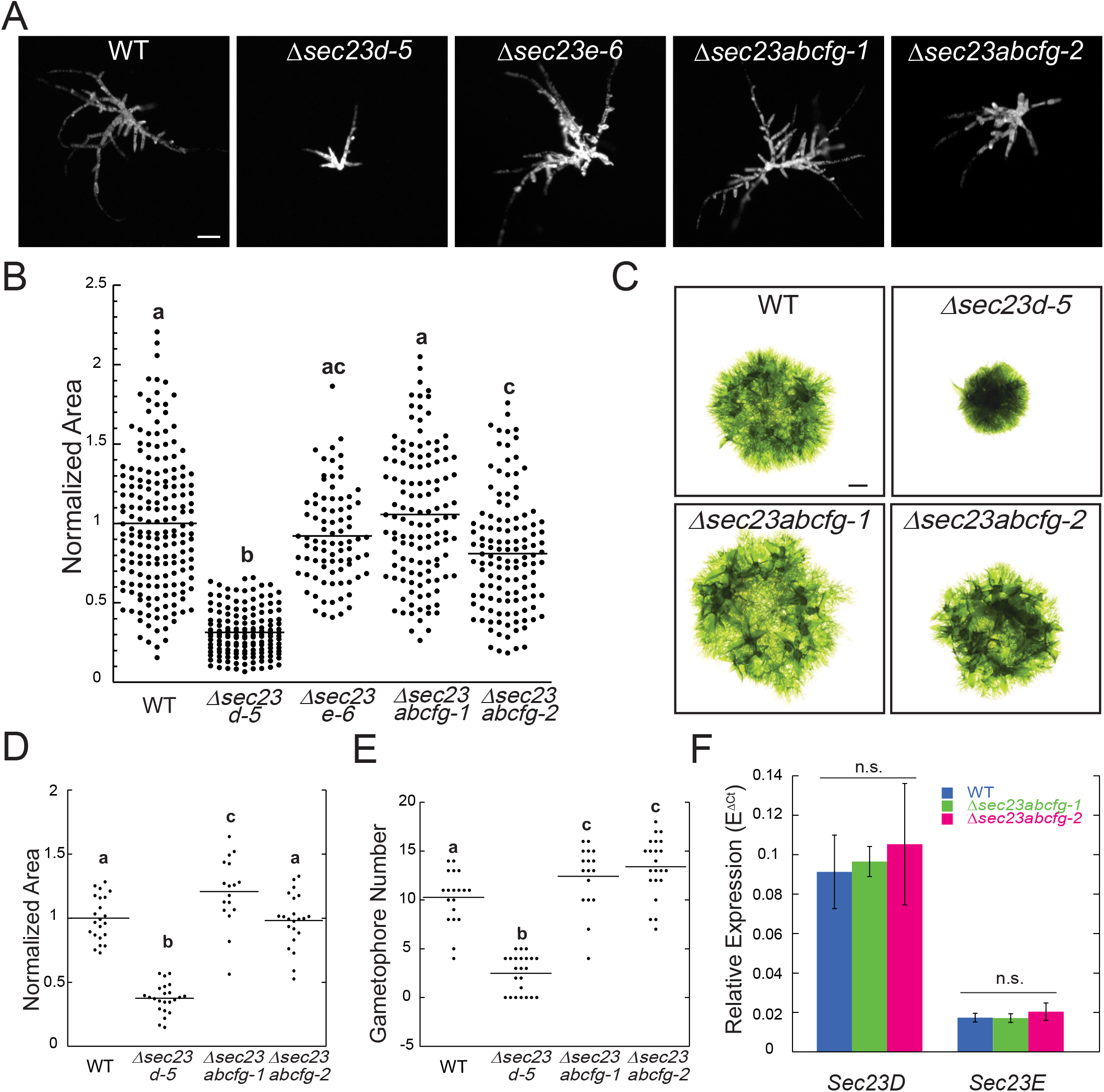
CRISPR-Cas9-mediated deletion of *Sec23* isoforms demonstrates that *Sec23D* is critical for protonemal growth. (A) Representative chlorophyll autofluorescence images of 7-day old plants with the indicated genotype regenerated from protoplasts. Scale bar, 100µm. (B) Quantification of plant area normalized to the wild type control. (C) Representative 3-week old moss plants with the indicated genotype. Scale bar, 1mm. (D) Quantification of the plant area of 3-week old moss plants normalized to that of the wild type control. (E) Number of gametophores in 3-week old moss plants. Letters in (B, D-E) indicate groups with significantly different means as determined by a one-way ANOVA with a Tukey *post hoc* test (α=0.05). (F) Relative expression of *Sec23D* and *E* genes normalized to UBIQUITIN10 in 8-day-old wild type and *Δsec23abcfg-1*, and *Δsec23abcfg-2* moss plants regenerated from protoplasts. N=3, Error bars are s.e.m. No significant difference was determined by a one-way ANOVA with a Tukey *post hoc* test (α=0.05).

Although it was possible to isolate many *Δsec23d* and *Δsec23e* single mutants, we were unable to obtain *Δsec23de* double mutants either by targeting *Sec23D* and *Sec23E* simultaneously or by targeting *Sec23D* in the *Δsec23e-6* single mutant background. This suggests that deleting both *Sec23D* and *Sec23E* is incompatible with protoplast regeneration, consistent with the severe growth defects observed by simultaneously silencing *Sec23D* and *Sec23E* using RNAi. Furthermore, we confirmed that none of the other *Sec23* genes were upregulated in *Δsec23d-5* plants (Fig S1C), suggesting that the phenotypic consequences observed were due specifically to loss of *Sec23D*.

Since the RNAi results suggested that all other *Sec23* subclasses were not required for protonemal growth, we sought to isolate a stable line lacking *Sec23A, Sec23B, Sec23C, Sec23F*, and *Sec23G* to determine if these genes contribute to other aspects of plant growth and development. We successfully isolated two independent quintuple mutant alleles *Δsec23abcfg-1* and *Δsec23abcfg-2*, in which *Sec23A, Sec23B, Sec23C, Sec23F*, and *Sec23G* contained lesions resulting in null alleles for each gene (Fig S2, Table S2). Most lesions were small deletions or insertions. However, *Δsec23abcfg-1* contained a three thousand base pair insertion in the *Sec23G* locus, still resulting in disruption of the *Sec23G* locus (Fig S2, Table S2). We characterized both quintuple mutants and found that while seven-day old *Δsec23abcfg-1* plants exhibited no detectable growth defects, *Δsec23abcfg-2* plants exhibited a mild growth defect (Fig 3B, C). However, three-week-old *Δsec23abcfg-1* plants were larger than wild type, and *Δsec23abcfg-2* plants were indistinguishable from wild type (Fig 3D, E). Interestingly, both *Δsec23abcfg-1* and *Δsec23abcfg-2* developed more gametophores than wild type (Fig 3E), suggesting that these five *Sec23* genes may negatively regulate gametophore development.

Even though the two quintuple mutant isolates, exhibited statistically significant differences at different times in development between each other and wild type, these were minor compared to the differences between wild type and all isolated *Δsec23d* null mutants. To investigate whether the observed differences might result from differential upregulation of *Sec23D* or *Sec23E* in the *Δsec23abcfg* mutants thereby compensating for the loss of *Sec23ABCFG* functions, we measured the expression levels of *Sec23D* and *Sec23E* transcripts in *Δsec23abcfg-1, Δsec23abcfg-2*, and wild type. In 8-day-old plants, we found that neither *Sec23D* nor *Sec23E* transcripts were elevated compared to expression in wild type (Fig 3F), indicating that minor differences observed between the two quintuple mutants were not readily explained by increased expression of either *Sec23D* or *E*. Taken together, the phenotypic analyses of stable *sec23* null plants suggest that *Sec23A, B, C, F* and *G* do not contribute substantially to establishment of juvenile moss tissues under laboratory growth conditions. In contrast, *Sec23D* and *E* are essential.

### Loss of Sec23D causes ER morphology defects and ER stress

As a predicted member of the COPII complex, Sec23D likely mediates ER-to-Golgi transport. However, it is unclear whether the *Δsec23d* growth defect is a result of a general impairment of ER-to-Golgi transport, or the lack of specific cargo sorted by Sec23D needed for protonemal growth. Studies in other organisms have shown that blocking COPII function alters ER morphology (Novick et al., 1980) and results in elevated levels of unfolded proteins in the ER, leading to activation of the ER unfolded protein response (Belden, 2001; Chung et al., 2016). To analyze Sec23D’s function in ER-to-Golgi transport, we investigated whether loss of Sec23D affects ER morphology and the ER unfolded protein response. We used CRISPR-Cas9 mediated genome editing to knock out Sec23D in a line where the ER is fluorescently labeled by targeting GFP to the ER lumen (Fig S1, Table S2). When imaged with a confocal microscope, *Δsec23d-B11* cells exhibited numerous ER dots (Fig 4A, red and cyan boxes), and 65% of protonemal filaments contained large ER aggregates (Fig 4A, green and yellow boxes). Imaging with a super resolution spinning disc confocal system allowed closer inspection of the aggregates (Fig. 4B). Smaller aggregates were clearly comprised of accumulated tubules, while large aggregates appeared to have tubules that wrapped around each other (Fig. 4B, Movie S1).

**Figure 4.**
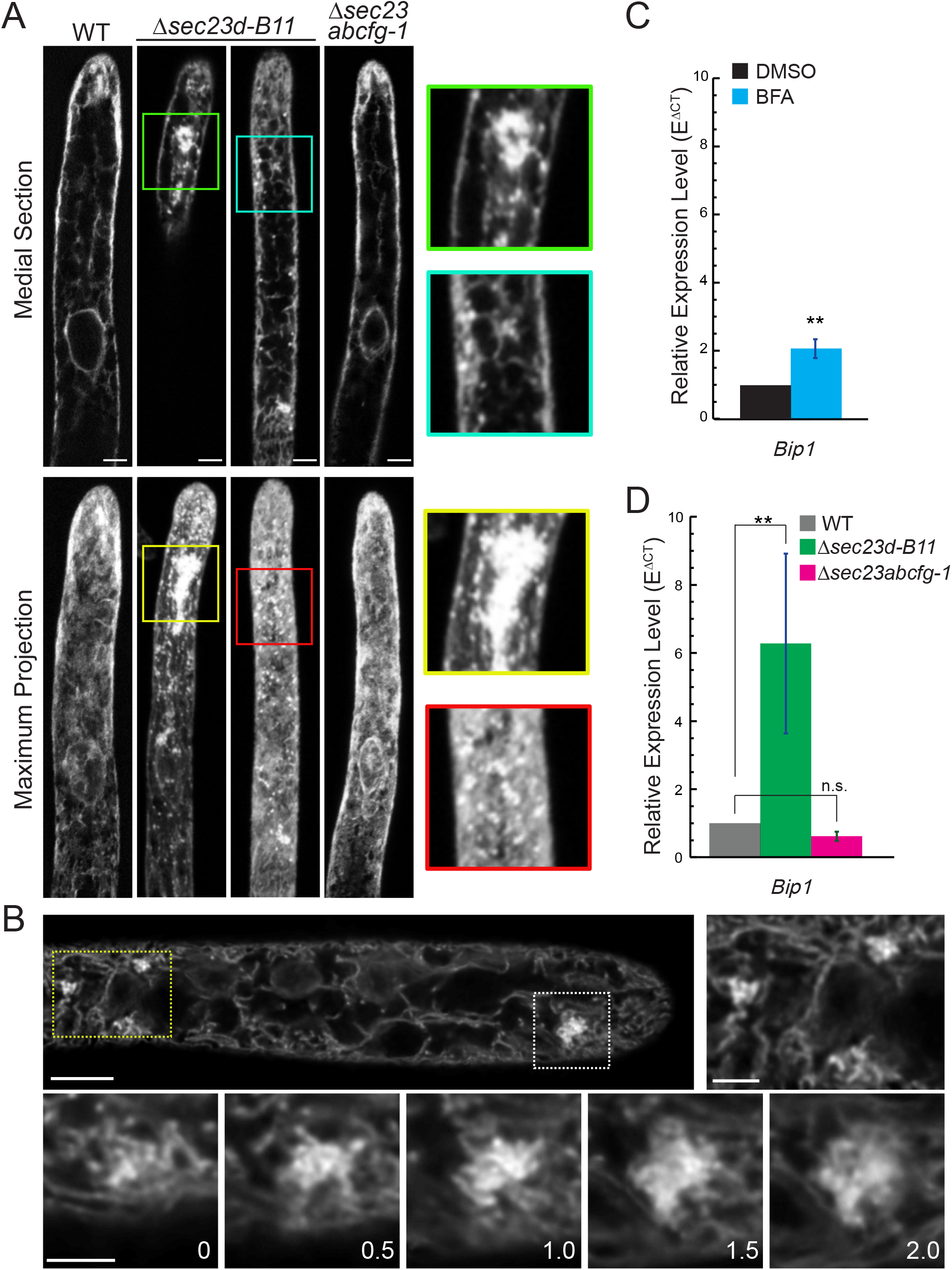
Loss of Sec23D causes ER morphology defects and ER stress. (A) Laser scanning confocal images of the ER labeled with GFP-KDEL in the indicated genotype. Colored boxes highlight large and small ER aggregates found in the *Δsec23d-B11* mutant. Scale bars, 5 µm. (B) Spinning disc confocal super resolution imaging of the ER labeled with GFP-KDEL in a *Δsec23d-B11* cell. Scale bar, 5 µm. Box outlined with yellow dotted line is enlarged to the right and the box outlined with a white dotted line is enlarged below with optical sections spaced every 0.5 µm shown for this region of the cell. Scale bars in boxed regions, 2 µm. Also see Movie S1. (C, D) Relative expression of *PpBip1* in 8-day old moss plants regenerated from protoplasts. N=3, Error bars are s.e.m. Asterisks denote t-probability is <0.01. n.s. denotes no significant difference. In (B) BFA-treated plants were normalized to DMSO-treated plants and in (C) null mutants were normalized to wild type plants.

If changes in ER morphology result from defective secretion at ER exit sites, then proteins may be inadvertently accumulating in the ER, resulting in upregulation of the unfolded protein response. To test this, we analyzed the expression of an ER heat shock protein (Bip), which is a well-known ER stress reporter in *Arabidopsis thaliana* (Noh et al., 2003; Srivastava et al., 2013; Maruyama et al., 2014; Cho and Kanehara, 2017). *P. patens* has two *Bip* paralogs, *Bip1* and *Bip2*. Since *Bip2* expression is low in protonemal tissue (Ortiz-Ramírez et al., 2017), we tested whether *Bip1* is affected under conditions that induce ER stress. Brefeldin A (BFA) inhibits protein secretion by disrupting ER-Golgi integrity (Nebenfuhr, 2002; Niu et al., 2005). We found that after a 24-hour treatment of BFA, expression of *Bip1* increased by about two-fold (Fig 4C), indicating that *Bip1* is a plausible ER stress reporter in *P. patens*. Interestingly, *Bip1* expression increased 5-fold in *Δsec23d* compared to in wild type (Fig 4D), demonstrating that *Δsec23d* suffers from severe ER stress. In contrast, *Δsec23abcfg-1*, which didn’t exhibit significant growth defects, had normal levels of *Bip1* expression (Fig 4D), and normal ER morphology (Fig 4A).

### Disrupting Sec23D results in reduced ER to Golgi trafficking and secretion to the plasma membrane

ER morphology defects and elevated ER stress in *Δsec23d* suggest that transport of secretory cargos might be blocked. We reasoned that proteins destined to the Golgi and the plasma membrane would accumulate in the ER in mutants with impaired ER to Golgi transport. To test this, we disrupted *Sec23D* in lines that either express YFP fused to the first 49 amino acids of the soybean α-1,2-mannosidase, a Golgi resident protein (YFP-MAN), (Nelson et al., 2007; Furt et al., 2012) or an mCherry fusion of a transmembrane protein (F-SNAP-mCherry) previously used to measure exocytosis (van Gisbergen et al., 2018) (Fig S1, Table S2). Since it would have been time-consuming to regenerate quintuple mutants in these two backgrounds, we instead transformed the YFP-MAN and F-SNAP-mCherry markers into *Δsec23abcfg-1* plants. We selected lines that had similar expression levels to control lines by measuring the total amount of fluorescence in cells.

In control YFP-MAN lines, YFP signal accumulated in the Golgi with very little fluorescence observed in the rest of the cell (Fig 5A). In contrast in *Δsec23d-F12*, while YFP fluorescence still accumulated in the Golgi, a large portion of the fluorescent signal that was presumably retained in the ER was observed diffusely throughout the cell (Fig 5A). To quantify ER to Golgi trafficking efficiency, we calculated the fluorescence intensity ratio of the signal in the Golgi divided by the signal in the cytoplasm. A high ratio value indicates efficient delivery to the Golgi, as observed in the control lines (Fig 5B). In contrast, *Δsec23d-F12* exhibited a significantly lower fluorescence intensity ratio (Fig 5B), indicating that loss of Sec23D impairs delivery of YFP-MAN to the Golgi. Imaging of *Δsec23abcfg-1* YFP-MAN and its control line revealed that the majority of the YFP signal localizes to the Golgi with the Golgi to cytoplasm ratios in *Δsec23abcfg-*1 indistinguishable from the control (Fig 5B), suggesting that trafficking of YFP-MAN to the Golgi is not affected in the quintuple mutant.

**Figure 5.**
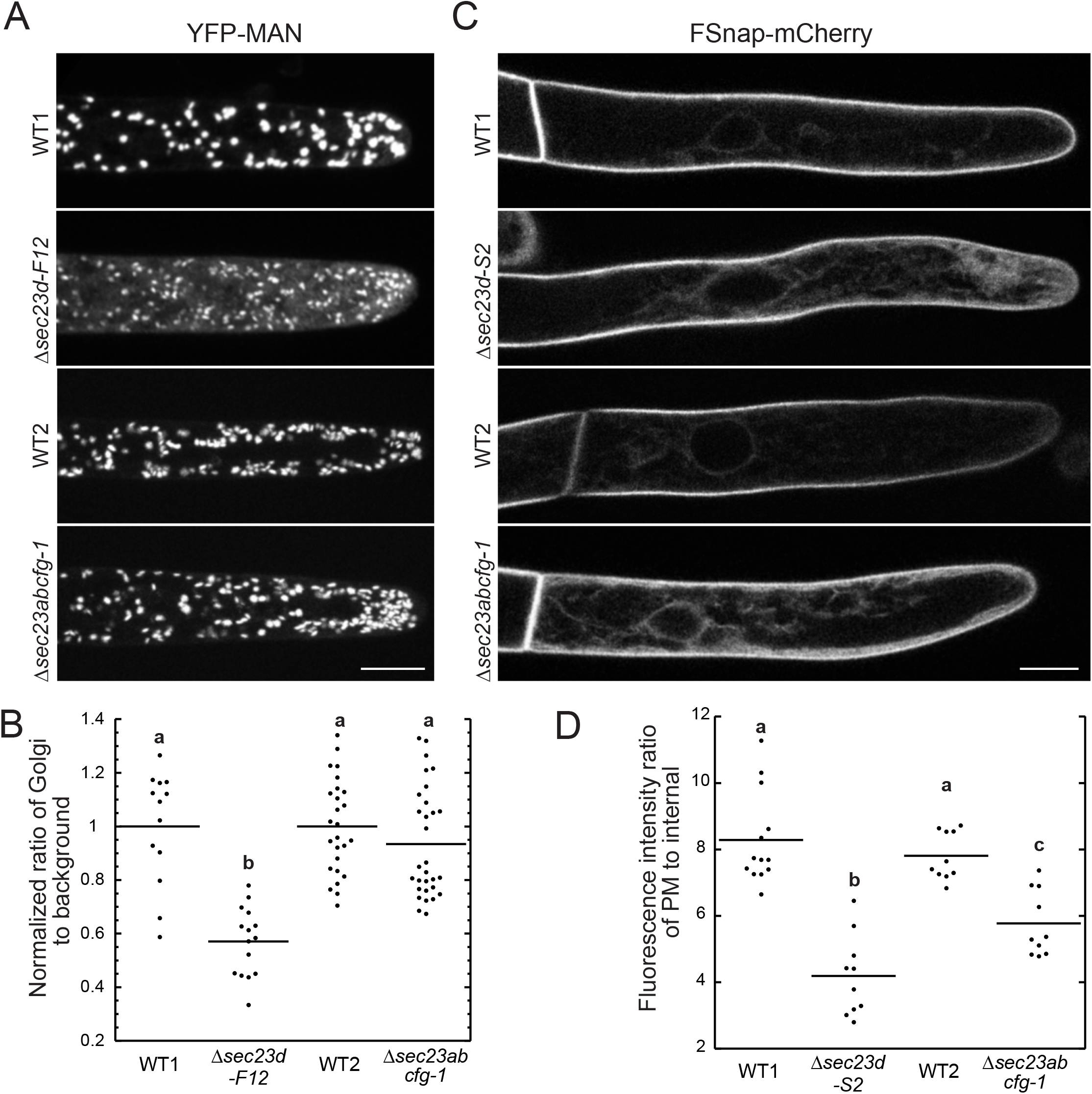
Deletion of *Sec23D* inhibits Golgi trafficking and secretion to the plasma membrane. (A) Representative maximum projections of confocal images of the indicated genotypes expressing a YFP-MAN marker. Scale bar, 10 µm. (B) Quantification of the fluorescence intensity ratio within the Golgi to the background. (C) Representative medial sections of confocal images of the indicated genotypes expressing a plasma membrane marker (FSnap-mCherry). Scale bar, 10 µm. (D) Quantification of the fluorescence intensity ratio of the plasma membrane to the intensity within the cell. Letters in (B, D) indicate groups with significantly different means as determined by a one-way ANOVA with a Tukey *post hoc* test (α=0.05).

Many proteins transported in COPII-coated vesicles ultimately reside at the plasma membrane, are secreted, or are trafficked to the vacuole. Without a fluorescent vacuolar cargo in hand, we instead investigated vacuolar morphology by staining the vacuole with MDY-64, a dye that stains the tonoplast in living plant cells (Scheuring et al., 2015). While we found that vacuolar morphology was similar between wild type, *Δsec23d-5* and *Δsec23abcfg-1* plants (Fig S3), we can not rule out that vacuolar protein sorting has not been affected.

To investigate trafficking to the plasma membrane, we used an engineered fluorescent protein cargo, F-SNAP-mCherry, previously used to analyze exocytosis in protonemata (van Gisbergen et al., 2018). In wild type cells, the F-SNAP-mCherry accumulated on the plasma membrane, with only a very small amount of fluorescence detected inside the cell (Fig 5C). In *Δsec23d-S2*, a large portion of the mCherry signal was retained in the cell, labeling the ER network (Fig 5C). To quantify secretion efficiency to the plasma membrane, we calculated the fluorescence intensity ratio of the plasma membrane to the intensity within the cell. We found that the ratio in *Δsec23d-S2* was significantly lower than that of its control line (Fig 5D), indicating that loss of Sec23D inhibits secretion of plasma membrane proteins.

Interestingly, we found that the F-SNAP-mCherry signal was also retained inside the cell in the *Δsec23abcfg-1* mutant, readily observing labeling of the ER network (Fig 5C). In addition, the fluorescence intensity ratio in *Δsec23abcfg-1* was lower in comparison to the ratio in its control line (Fig 5D). However, the ratio in *Δsec23abcfg-1* was still significantly higher than the ratio in *Δsec23d-S2*, suggesting that secretion of plasma membrane proteins is impaired in *Δsec23abcfg-1* but to a lower extent than in *Δsec23d-S2*. These data together with the lack of a growth defect in protonomata in the *Δsec23abcfg-1* mutant suggest that Sec23A, B, C, F, and G contribute to protein secretion particularly to the plasma membrane, but importantly their cargos are not essential for polarized growth in protonemata.

### Loss of Sec23D does not affect the cytoskeleton during tip growth

To investigate if defects in protein secretion, resulted in alterations to the cytoskeleton, critical for polarized expansion in protonemata, we imaged microtubules and actin in wild type and in cells carrying null mutations in *Sec23D*. Microtubules align along the length of the cell with their plus ends focused just below the cell apex (Hiwatashi et al., 2014), where a dynamic cluster of actin filaments forms predicting the site of cell expansion (Wu et al., 2018). Wild type and *Δsec23d-M4* cells expressing GFP-tubulin exhibit indistinguishable microtubule networks and both have a focal point of microtubules near the cell tip (Fig S4A, arrows). Using lifeact-mCherry to label the actin network, we found that both wild type and *Δsec23d-5* cells form an apical actin cluster that associates with the growing tip (Fig S4B). Time-lapse imaging revealed no discernable differences in the dynamics of the apical actin cluster between wild type and *Δsec23d-5* (Fig S4C, Movie S2). These data suggest that tip growth defects in cells lacking Sec23D are independent of the cytoskeleton.

### Sec23 isoforms localize to discrete puncta associated with the ER

If Sec23 is a bona fide COPII component, then we would expect that Sec23 isoforms should associate with the ER. To test this, we inserted sequences in the genome of a line expressing GFP in the ER lumen encoding for three tandem mRuby2 (hereafter, 3XmRuby) in frame with the coding sequences of the three mostly highly expressed protonemal *Sec23* genes: *Sec23B, Sec23D* and *Sec23G* (Fig 1B, Fig S5A, B). To ensure that tagging *Sec23D* did not affect its function, we measured plant area from 7-day old regenerated protoplasts and found that *Sec23D-3XmRuby* grew indistinguishably from the control lines (Fig S5C), suggesting that Sec23D-mRuby is functional. By analogy, we reasoned that tagging *Sec23B* and *G* would also be functional. To ensure we did not observe any possible dominant negative impacts as a result of the tagging, we performed growth assays of 7-day old plants regenerated from protoplasts and observed that *Sec23B-3XmRuby* was indistinguishable from the control and *Sec23G-mRuby* was slightly smaller (Fig S5C).

Imaging with confocal microscopy revealed that Sec23D, Sec23B and Sec23G localized to punctate structures throughout the cytoplasm (Fig 6A). All three proteins labeled puncta that were often proximal to the ER (Fig 6A). Co-localization measurements using Pearson’s Correlation Coefficient suggested that Sec23D, B and G puncta positively correlated with the ER. To ensure that this correlation was significant, we flipped the ER image both vertically and horizontally and repeated the correlation analysis and found that the correlation coefficients were significantly lower (Fig 6B). Interestingly, Sec23G-3XmRuby puncta were larger than the punctate structures found in Sec23D- and Sec23B-3XmRuby lines (Fig 6A, C). To quantify possible size differences, we imaged all three tagged Sec23 isoforms with a pixel size that maximizes the resolving capacity of the imaging system (Fig 6C, see Materials and Methods) and utilized a segmentation tool in Fiji that identifies particles in the images (Gilles et al., 2017). We discovered that Sec23G particles are significantly larger than Sec23B and D. Furthermore, Sec23D is slightly larger than Sec23B. These data suggest that Sec23 isoforms form distinct compartments on the ER.

**Figure 6.**
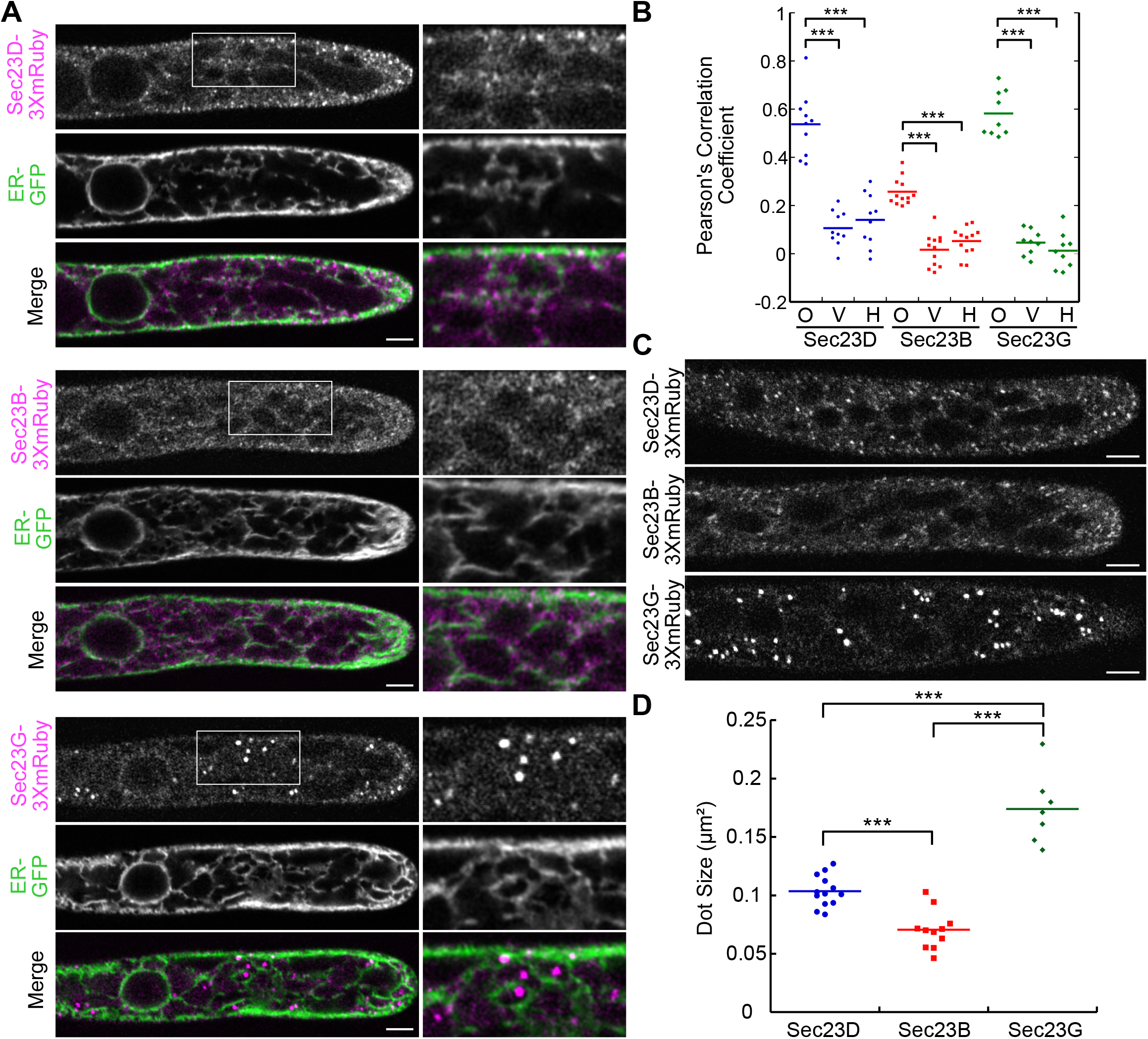
Sec23 isoforms localize to ER proximal structures with distinct sizes. (A) Representative confocal Z-sections of lines where the indicated Sec23 gene was endogenously tagged with 3XmRuby in a line that expresses GFP-KDEL labeling the ER lumen (ER-GFP). Scale bars, 5 µm. Boxed region in the mRuby channel is shown to the right of each image. (B) Pearson’s correlation coefficient measured between the Sec23 labeled lines and ER-GFP. O, original ER image; V, vertically flipped ER image; H, horizontally flipped ER image. Asterisks denote p<0.001as determined by a one-way ANOVA with a Tukey *post hoc* test (α=0.05) (B) Representative confocal Z-sections of *Sec23* genes endogenously tagged with 3XmRuby acquired with the optimal resolution of the confocal imaging system. Scale bars, 5 µm. (D) Quantification of average dot size in the Sec23 images as determined by the diAna tool for Fiji (Gilles et al., 2017). Asterisks denote p<0.001 as determined by a one-way ANOVA with a Tukey *post hoc* test (α=0.05).

### Sec23B and Sec23G localization is independent of Sec23D

To further probe the nature of the Sec23 particles, we analyzed the localization of Sec23B and Sec23G with respect to Sec23D. We used CRISPR-Cas9 mediated homology directed repair (HDR) to insert sequences encoding for mNeonGreen (hereafter, mNeon) in-frame with Sec23B or Sec23G in a Sec23D line endogenously tagged with a single mRuby2 (hereafter, mRuby) (Fig S6). As expected Sec23B-mNeon formed small puncta whereas Sec23G-mNeon formed larger puncta (Fig 7A, B). Imaging of Sec23B-mNeon and Sec23D-mRuby revealed the presence of a distribution of cytoplasmic puncta, some with only Sec23B-mNeon or only Sec23D-mRuby and some with both (Fig 7A). In contrast, Sec23G-mNeon and Sec23D-mRuby were largely non-overlapping (Fig 7B). To quantify the degree of overlap, we utilized the segmentation tool in Fiji that identifies puncta and then determines the number of segmented particles that physically overlap (Gilles et al., 2017). We found that 57% of Sec23B-mNeon co-localized with Sec23-mRuby. In contrast, only 4% of Sec23G-mNeon showed overlap with Sec23-mRuby (Fig 7C). In addition to identifying co-localizing particles, the tool also measures the center-to-center distance of overlapping particles (Gilles et al., 2017). Sec23G-mNeon and Sec23-mRuby had a significantly larger center-to-center distance as compared to Sec23B-mNeon and Sec23D-mRuby (Fig 7D), suggesting that any overlap between Sec23G and Sec23D is coincidental.

**Figure 7.**
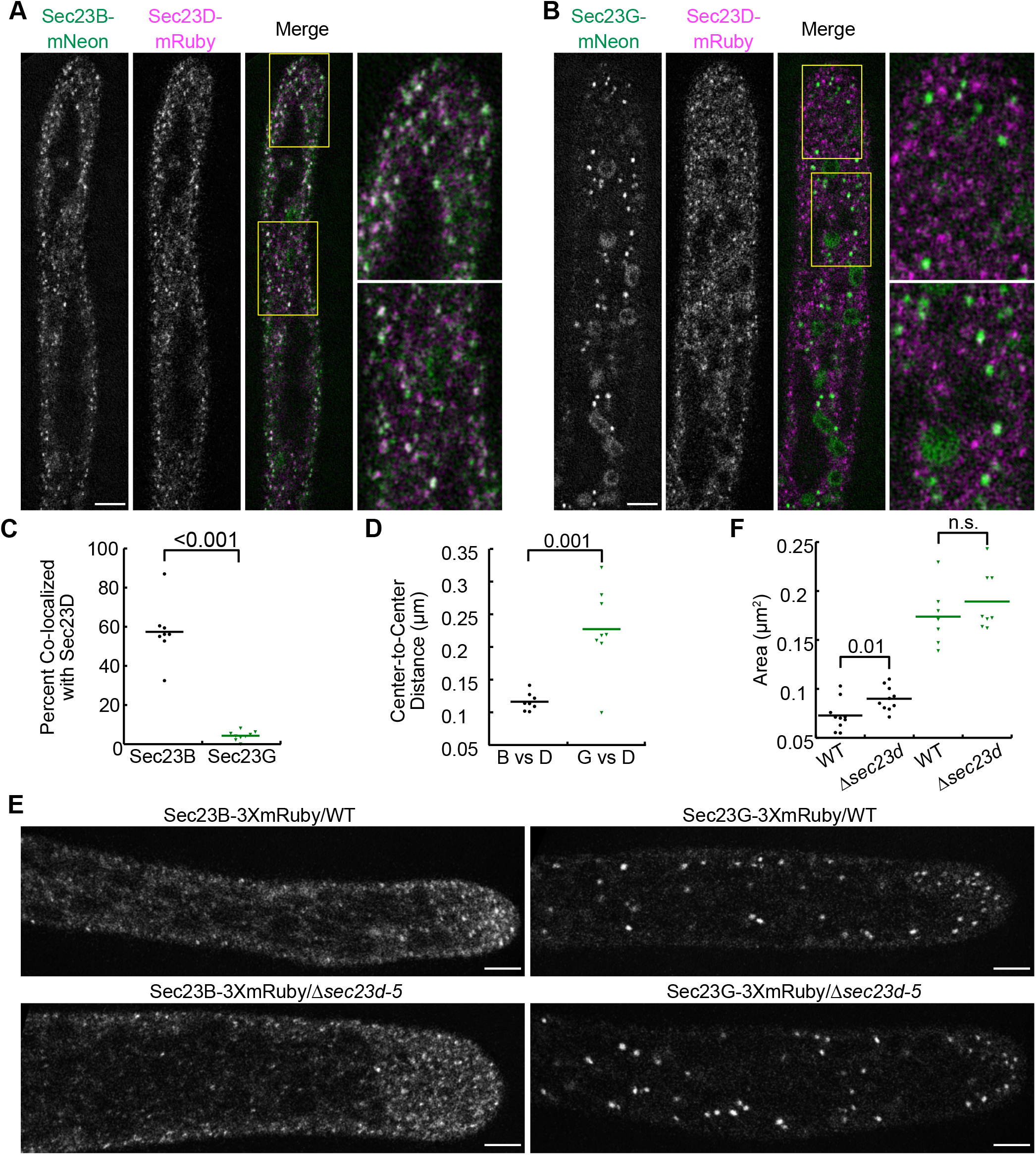
Sec23B and Sec23G form ER exit sites independent of Sec23D. Representative confocal Z-sections of Sec23D-mRuby lines where Sec23B (A) or Sec23G (B) were endogenously tagged with mNeon. Scale bar, 5 µm. Yellow boxed regions are enlarged to the right of the merged image where Sec23D-mRuby is false-colored magenta and in (A) Sec23B-mNeon is false-colored green and in (B) Sec23G-mNeon is false-colored green. (C) Percent of Sec23B-mNeon or Sec23G-mNeon dots that co-localized with Sec23D-mRuby and (D) center-to-center distance of co-localized particles as determined using the diAna tool for Fiji (Gilles et al., 2017). (E) Representative confocal Z-sections of wild type and *Δsec23d-5* cells with either Sec23B or Sec23G endogenously tagged with 3XmRuby. Scale bar, 5 µm. (F) Quantification of average dot size as determined by the diAna tool for Fiji (Gilles et al., 2017). Sec23B and Sec23G dot size in wild type is the same data presented in Figure 6D. Student t-test was performed for data in (C, D, and F). The t-probability is indicated for each data pair in each graph. n.s. denotes no significant difference.

While Sec23B and Sec23D exhibited a high degree of co-localization, COPII vesicles are significantly smaller than the limit of resolution of a light microscope. Therefore, it is unclear if a single vesicle contains both Sec23B and Sec23D or if vesicles form with a single Sec23 isoform. Given that Sec23G structures are distinct from Sec23D, and 40% of Sec23B and Sec23D puncta are not overlapping, we hypothesized that Sec23B, Sec23D and Sec23G form distinct ER exit sites. If this is the case, we would expect that Sec23B and Sec23G would be unaffected by loss of Sec23D. Indeed, we discovered that when we deleted Sec23D in the Sec23B-3XmRuby and Sec23G-3XmRuby lines, both Sec23B and Sec23G still localized to puncta (Fig 7E). We quantified the size of the puncta and found that Sec23G-3XmRuby was unaffected but Sec23B-3XmRuby were slightly larger in the absence of Sec23D (Fig 7F). However, given that the average Sec23B-3XmRuby particle size is essentially at the limit of resolution of our system, it is unclear how significant this increase may be. Nevertheless, these results demonstrate that even when ER morphology is affected due to loss of Sec23D, Sec23B and Sec23G are largely unaffected, suggesting that Sec23B and Sec23G ER exit sites are independent of Sec23D.

## Discussion

The large number of genes encoding components of COPII in plants together with functional studies in Arabidopsis (De Craene et al., 2014; Chung et al., 2016; Barlow and Dacks, 2018; Brandizzi, 2018; Faso et al., 2009; Qu et al., 2017; Tanaka et al., 2013) have led to the hypothesis that gene expansion underlies increased functional diversification required for complex secretory processes that pattern and build plant cells. Evidence suggests that functional diversification could stem from tissue/developmental specific expression (Hino et al., 2011; Lee et al.; Hanton et al., 2009) or from the formation of COPII complexes comprised of specific isoforms with unique functions (Zeng et al., 2015; Zeng et al., 2021). However, it remains unclear whether unique COPII complexes mediate transport of specific cargos or accumulate at some ER exit sites and not at others. Here, by analyzing the role of Sec23 and Sec24 isoforms in protonemal development, we have provided evidence for formation of distinct ER exit sites and the requirement of only Sec23D/E for transport of cargos uniquely required for tip growth.

Using RNAi, we showed that *Sec23* and *Sec24* isoforms differentially impact protonemal growth. Silencing of *Sec23D/E* or *Sec24C/D* similarly inhibit tip growth suggesting that *Sec23D/E* may interact with *Sec24C/D*. However, *Sec24F/G* also exhibited reduced growth. Thus, it is possible that a number of heterodimer pairs may form that are critical for polarized growth. Protein-protein interactions studies are needed to confirm if isoform specific heterodimers form in moss. If this is the case, it suggests that isoform specificity regulates COPII function similar to the finding in Arabidopsis that Sar1A and Sec23A form a unique interaction mediating functional specificity (Zeng et al., 2015).

Focusing on *Sec23*, where one of the Sec23 pairs, Sec23D/E, plays an essential role in protonemal growth, we found several lines of evidence suggesting that Sec23D functions as a bona fide COPII component. First, *Δsec23d* plants had abnormal ER morphology and elevated ER stress consistent with a block in cargo secretion from the ER. Second, trafficking to the Golgi and secretion to the plasma membrane was reduced in *Δsec23d*. Third, Sec23D localized to puncta near the ER. Due to differential expression levels, we found that Sec23D is more important for protonemal growth than Sec23E. However, when expressed at sufficient levels, Sec23E can rescue loss of Sec23D indicating that Sec23D and Sec23E are functionally redundant. The inability to recover a double mutant lacking both Sec23D and E suggests that together these two genes are essential for moss viability. Given that viability in moss is measured in the ability to generate protonemata, which is a tip-growing tissue, we reason that Sec23D and E are essential for tip growth. While *Δsec23d* protonemal cells grow significantly slower than wild type leading to smaller plants, we were unable to identify any defects in either the actin or microtubules cytoskeletons, suggesting that Sec23D contributes to polarized cell expansion independent of the cytoskeleton. In fact, tip growth in *Δsec23d* resembles the recently identified *P. patens Δsabre* mutant plants (Cheng & Bezanilla, 2021). SABRE associates with regions of the ER and its absence has no effect on the cytoskeleton but rather impacts ER morphology by accumulating aggregates that label with an ER lumenal marker (Cheng & Bezanilla, 2021). Taken together these data suggest that abrogated ER function impacts polarized growth without altering cytoskeletal organization and dynamics.

With respect to gametophore formation, *Δsec23d* develops gametophores late, but once gametophores form they are morphologically normal. This suggests that Sec23D function is specifically required during tip growth. In contrast, we found that the remaining five *Sec23* genes were dispensable for formation of both protonemata and gametophores. One possible explanation for this finding is that Sec23A, B, C, F and G have functionally diversified no longer playing a role in ER to Golgi transport. In support of this hypothesis, unlike *Δsec23d, Δsec23abcfg* plants did not exhibit ER morphology defects or increased ER stress characteristic of cargo backing up in the ER often observed with COPII loss of function mutants. However, we did observe secretion defects of plasma membrane cargo in *Δsec23abcfg* plants. Given that Sec23B and G localize proximal to the ER independent of Sec23D, it is possible that Sec23B and G contribute to COPII function via pathways that may potentially bypass the Golgi (Cheng et al., 2009; Zhang et al., 2011; Ding et al., 2012; Wang et al., 2017). Alternatively, Sec23B and G may also function in ER to Golgi trafficking but, in contrast to Sec23D, their cargos may be fewer and may not be essential for tip growth.

Interestingly all the mutant combinations that we generated formed morphologically normal gametophores. However, since we were unable to generate *Δsec23de* or *Δsec23abcefg* plants, we can not distinguish between Sec23E playing a critical role in gametophore formation versus Sec23D or Sec23E acting redundantly with Sec23A, B, C, F and G for COPII secretion required to shape gametophores. If, on the other hand, COPII is not required for gametophore formation, then future experiments conditionally removing Sec23D/E in *Δsec23abcfg* plants would be expected to form normal gametophores and would be an excellent tool to identify COPII-independent secretion pathways in plants, which are thought to play a role in transport of tonoplast resident proteins such as the vacuolar H^+^-ATPase, VHA-a3 (Viotti et al., 2013).

While Sec23B, Sec23D, and Sec23G formed puncta associated with the ER, these puncta were distinct. Sec23G formed large dots near the ER, whereas Sec23D and Sec23B were smaller. Sec23B and Sec23D puncta exhibited a high degree of overlap, suggesting that Sec23B and Sec23D may form vesicles in the same region of the ER. In contrast Sec23G largely did not overlap with Sec23D and by extension likely not Sec23B. Formation of larger domains on the ER suggests that Sec23G may be involved in the secretion of bigger cargo and provides evidence that moss has morphologically distinct ER exit sites, that can form, even when ER morphology is disrupted, independent of Sec23D.

Given that Sec23D contributes to ER to Golgi trafficking and secretion to the plasma membrane, while Sec23A, B, C, F and G contributes to plasma membrane secretion, it is plausible to suggest that tip growth requires intact ER to Golgi trafficking. In *Δsec23d* plants cargos essential for tip growth are either inefficiently carried from the ER to the Golgi or are not properly processed in the Golgi due to severe reductions in trafficking to the Golgi. Perhaps there is a larger burden for carbohydrates synthesized by enzymes in the Golgi to generate a tip growing cell wall as compared to a diffusely growing cell wall. To test this, future studies will focus on identifying cargos that have been differentially affected in *Δsec23d* versus *Δsec23abcfg* plants. Comparative mass spectrometry of isolated microsomes and plasma membrane extracts in these mutant backgrounds could be very informative. In addition, quantitatively measuring secretion of known Golgi resident carbohydrate biosynthetic enzymes and plasma membrane complexes such as cellulose and callose synthases in these mutants could help to narrow down cargo specificity.

Taking advantage of the facile molecular genetic manipulation and imaging in moss protonemal development, our study has shown that the expanded *Sec23* gene family in moss is multi-faceted. We found evidence that the *Sec23* gene family exhibits aspects of functional redundancy, differential gene expression, as well as isoform specificity for ER exit site formation, ER to Golgi trafficking, and importantly transport of essential cargos required for tip growth. Building upon the tools developed in this study, future studies will be able to characterize the remaining COPII components, thereby providing a complete picture of how gene expansions in COPII have been deployed.

## Materials and Methods

### Expression analyses

To quantify expression of the *Sec23* and *Sec24* genes family members (Table S3), as well as *Bip1* (Pp3c10_17310), we isolated total RNA from 8-day old plants (various genotypes, as needed for the particular experiment) regenerated from protoplasts using the RNeasy Plant mini prep kit (Qiagen). cDNA was prepared from 1µg of total RNA using SuperScriptIII reverse transcriptase kit, following the manufacturer’s recommendation (Invitrogen). qPCR assays were conducted with Luna Universal qPCR Master Mix (New England Biolabs). Primers for qRT-PCR analyses are listed in Table S4. After confirming that amplification efficiencies (calculated from standard curves) were similar for each set of primers (Table S4), relative expression levels were calculated, and expression of Ubiquitin 10 was used for normalization. All experiments had three technical replicates and three biological replicates.

### Constructs

The Sec23 and Sec24 RNAi constructs were generated by PCR amplification of either the coding sequence or 5’ and 3’ untranslated region (UTR) of *Sec23* or *Sec24* from *P. patens* cDNA using primers indicated in Table S4. PCR fragments were cloned into pENTR-D-TOPO (Invitrogen) following the manufacturer’s recommendations and the resulting vectors were sequenced. The Sec23ACD-RNAi and Sec23 UTR-RNAi constructs were generated by stitching together multiple PCR fragments and then directionally cloned into pENTR-D-TOPO as previously described (Vidali et al., 2007). LR clonase (Invitrogen) reactions were used to transfer either the coding sequence or 5’ and 3’ UTR sequences the RNAi vector pUGGi (Bezanilla et al., 2005) generating the Sec23 UTRi or Sec24-CDSi constructs, respectively. Restriction enzyme digestion was used to verify these constructs.

Expression constructs were generated by PCR amplifying *Sec23D* and *Sec23E* coding sequences from *P. patens* cDNA using a 5’ CACC sequence on the specific primers for oriented cloning into the pENTR-D-TOPO vector (Invitrogen). LR clonase (Invitrogen) reactions were used to transfer these coding sequence into pTH-Ubi-Gate vectors (Vidali et al., 2007).

*Sec23* tagging constructs were generated by PCR amplifying genomic sequences upstream and downstream of the stop codon for *Sec23D, Sec23B*, and *Sec23G* using the primers in Table S4. For *Sec23D*, we used homologous recombination. The distance between the 5’ and 3’ homology arms was 500bp, which when inserted correctly the tagging construct replaces these 500bp with sequences encoding for the fluorescent protein and an antibiotic resistance cassette (Fig S5, Fig S6). Using BP clonase (Invitrogen), homology arms were transferred to vectors, generating homology arm entry clones. Four-fragment recombination reactions were used to assemble the homology arms, fluorescent protein sequence, and the antibiotic resistance cassette into pGEM-gate (Vidali et al., 2009). For *Sec23B* and *Sec23G* tagging, we used CRISPR Cas9-mediated HDR. 5’ and 3’ homology arms upstream and downstream of the stop codon, with a distance between these two arms of 38bp for *Sec23B* and 131bp for *Sec23G* were PCR amplified from genomic DNA (Fig S5, Fig S6). Using BP clonase (Invitrogen), homology arms were transferred to vectors, generating homology arm entry clones, as described in (Mallett et al., 2019). Three-fragment recombination reactions were used to assemble the homology arms and the fluorescent protein sequence into pGEM-gate. We chose the CRISPR-Cas9 protospacer close to the stop codon.

Protospacers for CRISPR-Cas9 mediated mutagenesis and HDR were designed with the CRISPOR online software (crispor.tefor.net) (Haeussler et al., 2016). Protospacers were selected with high specificity scores and low off-target frequency. Specific overhangs sequences were added to facilitate downstream cloning. Protospacers were synthesized as oligonucleotides (Table S4) and annealed as described (Mallett et al., 2019). Annealed protospacer fragments were transferred to vectors using Instant Sticky-End Ligation Master Mix (New England Biolabs). *Sec23B* and *G* protospacers designed to tag these genes via HDR as well as *Sec23A, D* and *E* protospacers designed to knock out these genes were ligated into pENTR-PpU6-sgRNA-L1L2 (Mallett et al., 2019). *Sec23B* and *F* protospacers designed to knock out these genes were ligated into pENTR-PpU6-sgRNA-L1L5 (Mallett et al., 2019). Sec23C and G protospacers designed to knock out these genes were ligated into pENTR-PpU6-sgRNA-R5L2 (Mallett et al., 2019). LR clonase (Invitrogen) was used to assemble the protospacer containing vectors into pZeo-Cas9-Gate, pMH-Cas9-Gate, or pMK-Cas9-Gate vectors (Mallett et al., 2019), as needed. Two-fragment recombination reactions were used to assemble dual targeting mutagenesis vectors for *Sec23B* and *C*, and for *Sec23F* and *G*. Restriction enzyme digestion and sequencing was used to verify these constructs.

### Tissue propagation, transformation, and genotyping

All moss lines were propagated weekly by moderate homogenization in water and pipetting onto a permeable cellophane covered solid media, as described previously (Wu and Bezanilla, 2014). Tissue was grown at room temperature in Percival growth chambers, under 85 µmol_photons_/m^2^s light with long-day conditions. PEG-mediated transformations were performed as previously described to generate stable lines (Wu and Bezanilla, 2014), CRISPR-Cas9 genome edited lines (Mallett et al., 2019) and perform transient RNAi (Vidali et al., 2007).

To genotype mutants generated by CRISPR-Cas9 mutagenesis, we extracted DNA from plants that were 3-4 weeks old (0.5-1cm in diameter) using the protocol described in (Augustine et al., 2011). For editing experiments, we used primers (Table S4) surrounding the expected Cas9 cleavage site (∼300-400 bp on each side). T7 endonuclease assay was used to screen candidate mutants as described (Mallett et al., 2019). For homology-directed repair tagging and homologous recombination tagging, we used primers (Table S4) outside of the homology region to avoid amplification of residual DNA donor template.

### Imaging and morphometric analysis of growth assays

For transient RNAi, complementation analyses, and growth assays of stable lines, plants regenerated from protoplasts were photographed seven days after protoplasting. For transient RNAi and complementation analyses plants lacking nuclear GFP signal were imaged because these are the plants that are actively silencing as described previously (Bezanilla et al., 2005; Vidali et al., 2007). For growth assays of stable lines any regenerated plant was imaged. Images were acquired with a 1X objective using a stereomicroscope (Leica MZ16FA or Nikon SMZ25) equipped with a CCD camera (Leica DF300FX or Nikon digital sight DS-Fi2). Chlorophyll fluorescence and any nuclear GFP signal were acquired simultaneously using filters (480/40, dichroic 505 (Leica) or dichroic 510 (Nikon), emission 510 long pass). Exposure settings were maintained constant throughout an experiment. 3-week old plants regenerated from protoplasts were imaged with white light on a Nikon SMZ25 stereomicroscope equipped with a Nikon digital sight DS-Fi2 color camera.

To quantify plant area, individual plants were cropped from images and the red channel, representing chlorophyll fluorescence of the plant, was separated from the RGB image. Threshold settings were set manually for all plants within an experiment. To quantify brightfield images of 3-week-old plants, the blue channel, representing chlorophyll fluorescence of the plant, was separated from the RGB image and thresholded. Total area was calculated from the largest thresholded object in the selected window. All images analysis was done using macros (Galotto et al., 2019) written for ImageJ. Analysis of variation (ANOVA) for multiple comparisons was performed with Kaleidagraph (Synergy) using the Tukey HSD post hoc tests. The alpha for statistical significance was set to 0.05.

### Laser scanning confocal microscope imaging and quantification

For imaging Sec23B/D/G localization and dynamics, 5 to 8-day old plants regenerated from protoplasts were placed onto an agar pad in Hoagland’s buffer (4mM KNO_3_, 2mM KH_2_PO_4_, 1mM Ca(NO_3_)_2_, 89µM Fe citrate, 300µM MgSO_4_, 9.93µM H_3_BO_3_, 220nM CuSO_4_, 1.966µM MnCl_2_, 231nM CoCl_2_, 191nM ZnSO4, 169nM KI, 103nM Na_2_MoO_4_, and 1% sucrose), covered by a glass cover slip and sealed with VALAP (1:1:1 parts of Vaseline, lanoline, and paraffin). For imaging ER morphology, Golgi resident protein secretion, F-SNAP-mCherry secretion, we used microfluidic imaging chambers (Bascom et al., 2016). Ground protonemal tissue was gently pipetted into the central part of the device followed by an infusion of Hoagland’s medium. Then the chamber was submerged Hoagland’s medium, and placed under constant 85 µmol_photons_/m^2^s light.

Images were acquired on a Nikon A1R confocal microscope system with a 1.49 NA 60X oil immersion objective (Nikon) at room temperature. Image acquisition was controlled by NIS-Element AR 4.1 software (Nikon). Laser illumination at 488 nm was used for exciting mNeon, YFP, GFP and chlorophyll autofluorescence; 561 nm for mRuby, mCherry. Emission filters were 525/50 nm for mNeon, YFP, and GFP; 595/50 nm for mRuby and mCherry.

Super resolution spinning disc confocal imaging was performed with a Nikon CSUW1-SoRA system. Images were acquired with a 60x Plan Apo 1.40NA objective with the 2.8x mag lens. Laser illumination at 488 nm was used for exciting GFP, and the Emission filter were 525/50. Images were denoised using the denoise.ai machine learning algorithm and deconvolved using the Richardson-Lucy algorithm within the NIS-Elements v5.21 software.

For quantification of trafficking to the Golgi, confocal Z-stacks of individual cells were converted to a maximum intensity projection. For quantification of secretion to the plasma membrane, the medial focal plane was selected for analysis. YFP (YFP-MAN) and mCherry (FSnap-mCherry) fluorescence intensities with the same expression pattern were normalized using enhanced contrast in Fiji. Images were cropped to isolate regions only within the cell. Then, using a minimum threshold of 20,000-22,000 it was possible to isolate and measure the fluorescence intensity of the Golgi structures (Golgi intensity) or the plasma membrane with analyze particles in Fiji. The fluorescence intensity of the background was similarly measured using the same threshold cut-off for each image but unchecking dark background, thereby selecting all regions below the threshold cut off. These values were used to calculate the ratio of Golgi/plasma membrane intensity to background intensity. To quantify the Pearson’s Correlation Coefficient, we analyzed single focal planes with the Coloc2 plugin in Fiji comparing the original images as well as horizontally and vertically flipped ER-GFP images. To quantify the size and co-localization of Sec23B, D and G dots, we analyzed single focal planes and used the diAna tool for ImageJ, which uses object-based identification for distance and co-localization measurements (Gilles et al., 2017) and was recently used to define specific subdomains of the trans Golgi network in plant cells (Heinze et al., 2020). All images were acquired with a pixel size of 0.07 µm and processed by subtracting background, enhancing contrast with normalization, and smoothing in Fiji prior to performing segmentation with diAna. For Sec23G-mNeon, chloroplast autofluorescence was collected in the far-red channel and subtracted from the green channel. To segment the dots in the image, we used the iterative option with a minimum threshold of 25,000-27,500 and a step value of 20. After segmentation, the diAna analyse module was used to measure the size of all segmented regions of an image, as well as to determine the percent of co-localizing particles. For particles that did overlap, we also extracted the center-to-center distance of those overlapping particles. For all data, analysis of variation (ANOVA) for multiple comparisons was performed with Kaleidagraph (Synergy) using the Tukey HSD post hoc tests. The alpha for statistical significance was set to 0.05. For pairwise comparisons, a student-T test was used using Kaleidagraph (Synergy).

## Author contributions

M.C. and M.B. designed the research. M.C., S.W., J.E.O. and S.E.R performed the research. M.C., S.W. and M.B. wrote the manuscript.

## Acknowledgements

We thank members of the Bezanilla lab for careful reading of the manuscript. M.C. received support from the Plant Biology Graduate Program at the University of Massachusetts, Amherst. This work was supported by Dartmouth College and grants from the National Science Foundation (MCB-1330171 and MCB-1715785 to M.B.).

