## Supplemental Material for "COPII Sec23 proteins form isoform-specific ER exit sites with differential effects on polarized growth"

### Supplemental Information

#### *Supplemental Figure Legends*

**Figure S1. CRISPR-Cas9 mediate edited of *Sec23D* and *Sec23E*.** (A) Gene models are shown with exons indicated by boxes and introns by thin black lines. Coding and untranslated regions are denoted by thick and thin boxes, respectively. Arrows indicate the protospacer target site. Sequences in red denote the protospacers, underlined sequence denote the PAM motif. All mutants have indicated mutations, resulting in early translational stop; full-length Sec23D and Sec23E protein are 783 amino acids in length. (B) Quantification of plant area is based on the area of the chlorophyll autofluorescence and is presented normalized to that of the wild type control. Letters indicate groups with significantly different means as determined by a one-way ANOVA with a Tukey *post hoc* test ($\alpha$=0.05). (C) Relative expression of *Sec23* genes in 8-day old moss plants regenerated from protoplasts (normalized to the expression of UBIQUITIN10). N =3 Error bars are s.e.m.

**Figure S2. CRISPR-Cas9 mediate edited of *Sec23A, B, C, F* and *G*.** (A) Gene models are shown with exons indicated by boxes and introns by thin black lines. Coding and untranslated regions are denoted by thick and thin boxes, respectively. Arrows indicate the protospacer target site. Sequences in red denote the protospacers, underlined sequence denote the PAM motif. All mutants have indicated mutations, resulting in early translational stop; full-length Sec23A, B, C, F and G are 770, 779, 594, 816, and 934 amino acids in length, respectively.

**Figure S3. Vacuolar morphology is unaffected in *Sec23* mutant plants.** Representative confocal Z-sections of cells of the indicated genotype stained with MDY-64. Scale bars, 5 µm.

**Figure S4. Cytoskeleton is unaffected in *Δsec23d* plants.** Representative maximum projections of confocal Z stacks in the medial region of wild type and *Δsec23d* cells expressing GFP-tubulin (A) or Lifeact-mCherry (B). Scale bars, 10 µm in (A) and 5 µm in

(B). (C) Images from time-lapse acquisition of a single focal plane from wild type and Δsec23d cells expressing Lifeact-mCherry. Scale bars, 5 μm. Also see, Movie S2.

**Figure S5. Molecular characterization of endogenous tagging of *Sec23D, B* and *G* with 3XmRuby.** (A) Diagrams illustrate the result of homologous recombination (top) and HDR (middle and bottom) mediated insertion of 3XmRuby sequences in the genomic locus of each of the *Sec23* genes. Coding exons are indicated by thick boxes and untranslated exons are indicated by thin boxes. Thin lines indicate intronic regions and thin dotted lines are intronic regions downstream of the gene. Inserted sequences are denoted by thick colored boxes (magenta, 3XmRuby; yellow, hygromycin resistance cassette). The dashed vertical lines indicate the junction between the knock-in construct and upstream and downstream genomic sequences. Small arrows above the diagrams represent primers used for genotyping. Scale bar is 0.5 kb. (B) PCR products obtained with the indicated primer pairs using the template DNA isolated from the indicated moss line were separated on an agarose gel and stained with ethidium bromide. Molecular weight is indicated in kb. Predicted sizes for correct products are indicated in (A). (C) Quantification of plant area is based on the area of the chlorophyll autofluorescence and is presented normalized to that of the wild type ER-GFP control. Statistics were calculated with a one-way ANOVA with a Tukey *post hoc* test ($\alpha$=0.05).

**Figure S6. Molecular characterization of endogenous tagging of *Sec23 B* and *G* with mNeon and Sec23D with mRuby.** (A) Diagrams illustrate the result of homologous recombination-mediated insertion of mRuby into the Sec23D locus (top) and HDR-mediated insertion of mNeon into the *Sec23B* (middle) and *Sec23G* (bottom) loci. Coding exons are indicated by thick boxes and untranslated exons are indicated by thin boxes. Thin lines indicate intronic regions and thin dotted lines are intronic regions downstream of the gene. Inserted sequences are denoted by thick colored boxes (magenta, mRuby; yellow, hygromycin resistance cassette; light green, mNeon). The dashed vertical lines indicate the junction between the knock-in construct and upstream and downstream genomic sequences. Small arrows above the diagrams represent primers used for genotyping. Scale bar is 0.5 kb. (B) PCR products obtained with the indicated primer pairs using the template

DNA isolated from the indicated moss line were separated on an agarose gel and stained with ethidium bromide. Molecular weight is indicated in kb. Predicted sizes for correct products are indicated in (A).

**Supplemental Movie Legends**

**Movie S1. ER tubules aggregate in *Δsec23d* mutants.** Z-sections every 0.5 µm from a super resolution spinning disc confocal Z-stack, see Figure 4B. Movie plays at 5 fps.

**Movie S2. Actin dynamics are not altered in Δsec23d cells.** Single focal plane images from a time-lapse acquisition of wild type and Δsec23d cells labeled with lifeact-mCherry. Movie plays at 5 fps.

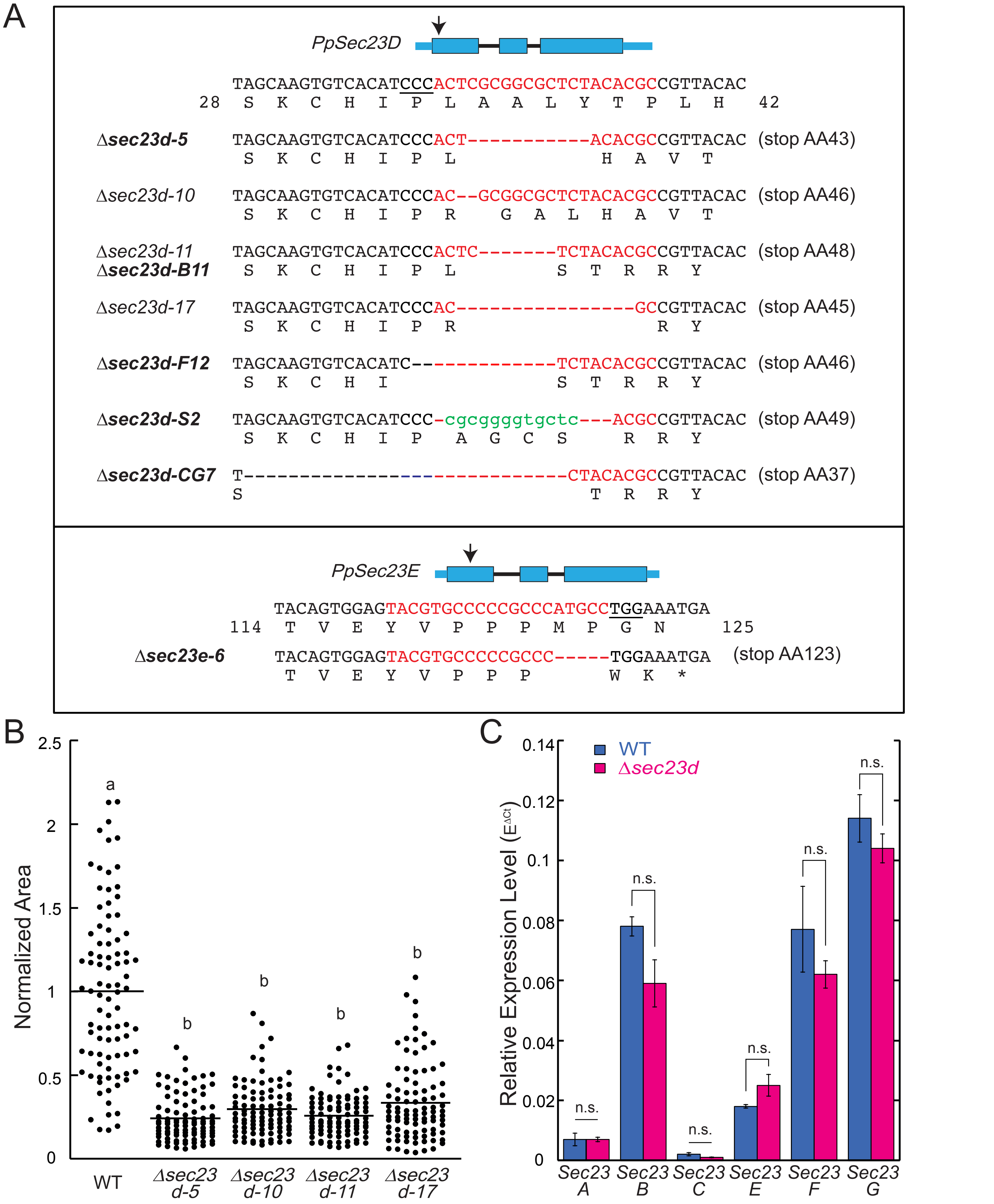


**Figure S1. CRISPR-Cas9 mediate edited of *Sec23D* and *Sec23E*.** (A) Gene models are shown with exons indicated by boxes and introns by thin black lines. Coding and untranslated regions are denoted by thick and thin boxes, respectively. Arrows indicate the protospacer target site. Sequences in red denote the protospacers, underlined sequence denote the PAM motif. All mutants have indicated mutations, resulting in early translational stop; full-length Sec23D and Sec23E protein are 783 amino acids in length. (B) Quantification of plant area is based on the area of the chlorophyll autofluorescence and is presented normalized to that of the wild type control. Letters indicate groups with significantly different means as determined by a one-way ANOVA with a Tukey *post hoc* test (α=0.05). (C) Relative expression of *Sec23* genes in 8-day old moss plants regenerated from protoplasts (normalized to the expression of UBIQUITIN10). N =3 Error bars are s.e.m.

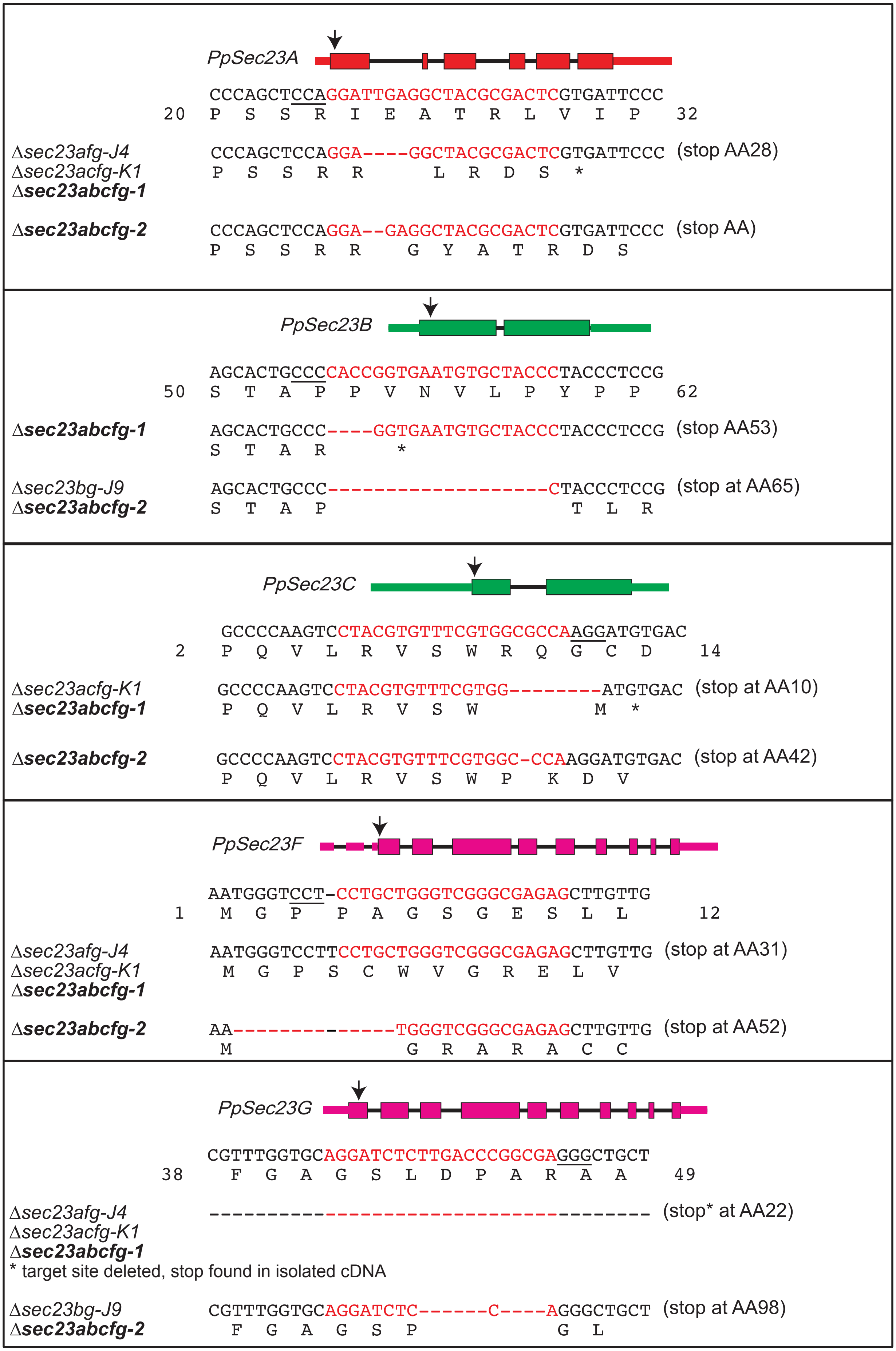


**Figure S2. CRISPR-Cas9 mediate edited of *Sec23A, B, C, F* and *G*.** (A) Gene models are shown with exons indicated by boxes and introns by thin black lines. Coding and untranslated regions are denoted by thick and thin boxes, respectively. Arrows indicate the protospacer target site. Sequences in red denote the protospacers, underlined sequence denote the PAM motif. All mutants have indicated mutations, resulting in early translational stop; full-length Sec23A, B, C, F and G are 770, 779, 594, 816, and 934 amino acids in length, respectively.

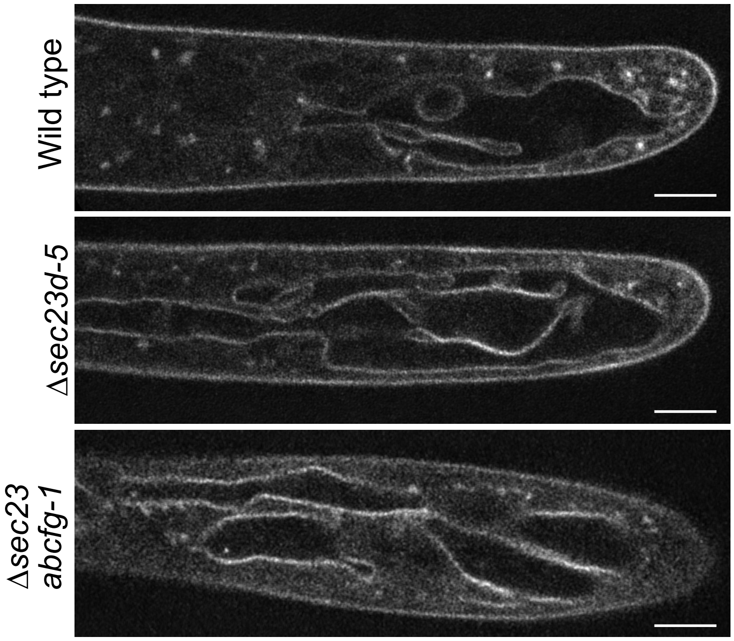


**Figure S3. Vacuolar morphology is unaffected in *Sec23* mutant plants.** Representative confocal Z-sections of cells of the indicated genotype stained with MDY-64. Scale bars, 5 μm.

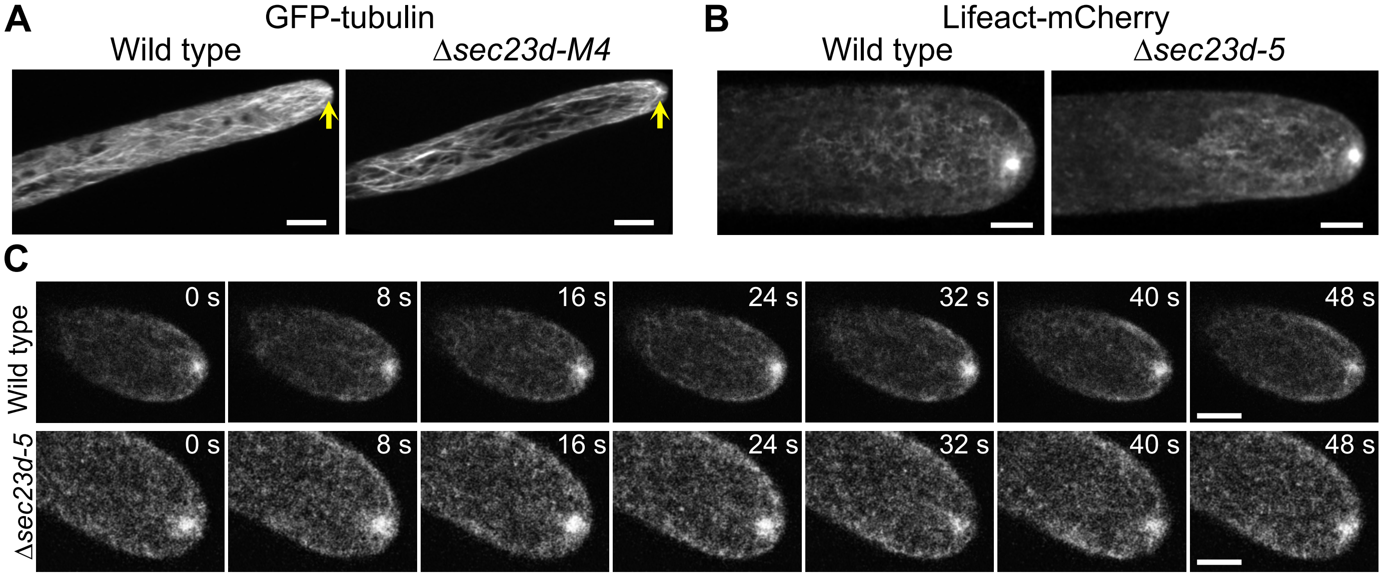


**Figure S4. Cytoskeleton is unaffected in Δ*sec23d* plants.** Representative maximum projections of confocal Z stacks in the medial region of wild type and Δ*sec23d* cells expressing GFP-tubulin (A) or Lifeact-mCherry (B). Scale bars, 10 µm in (A) and 5 µm in (B). (C) Images from time-lapse acquisition of a single focal plane from wild type and Δsec23d cells expressing Lifeact-mCherry. Scale bars, 5 µm. Also see, Movie S2.

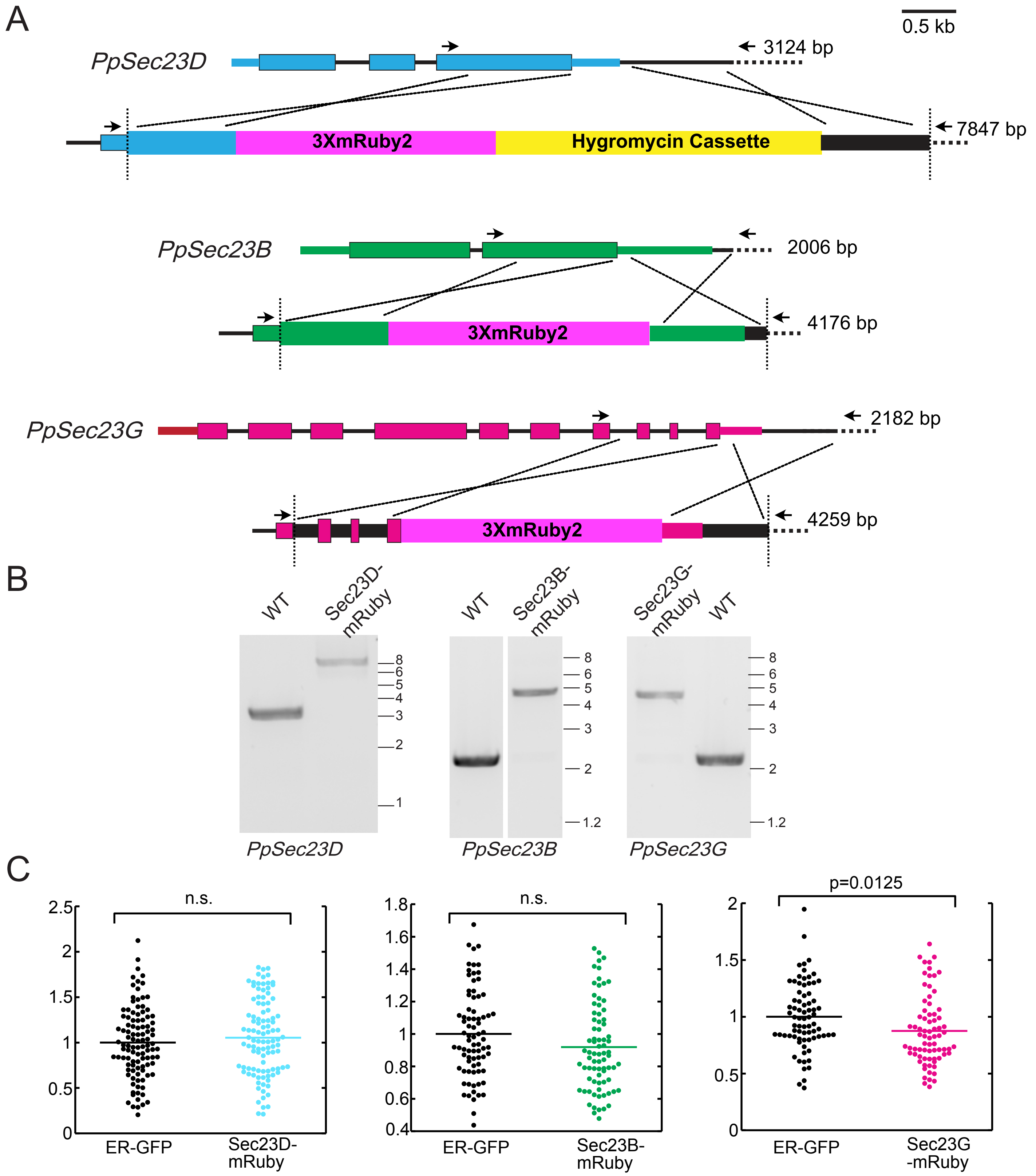


**Figure S5. Molecular characterization of endogenous tagging of *Sec23D, B* and *G* with 3XmRuby.** (A) Diagrams illustrate the result of homologous recombination (top) and HDR (middle and bottom) mediated insertion of 3XmRuby sequences in the genomic locus of each of the *Sec23* genes. Coding exons are indicated by thick boxes and untranslated exons are indicated by thin boxes. Thin lines indicate intronic regions and thin dotted lines are intronic regions downstream of the gene. Inserted sequences are denoted by thick colored boxes (magenta, 3XmRuby; yellow, hygromycin resistance cassette). The dashed vertical lines indicate the junction between the knock-in construct and upstream and downstream genomic sequences. Small arrows above the diagrams represent primers used for genotyping. Scale bar is 0.5 kb. (B) PCR products obtained with the indicated primer pairs using the template DNA isolated from the indicated moss line were separated on an agarose gel and stained with ethidium bromide. Molecular weight is indicated in kb. Predicted sizes for correct products are indicated in (A). (C) Quantification of plant area is based on the area of the chlorophyll autofluorescence and is presented normalized to that of the wild type ER-GFP control. Statistics were calculated with a one-way ANOVA with a Tukey *post hoc* test (α=0.05).

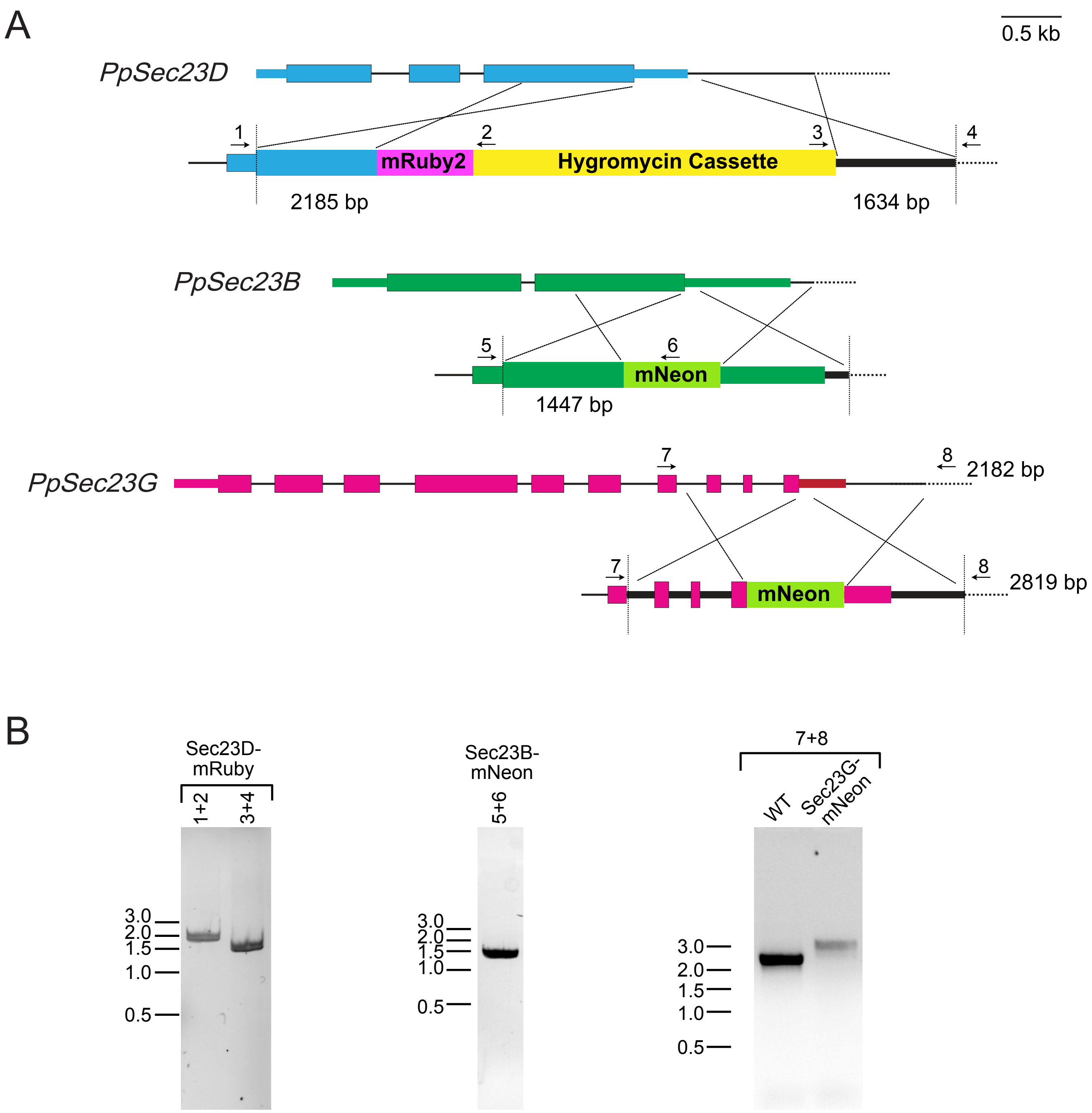


**Figure S6. Molecular characterization of endogenous tagging of *Sec23 B* and *G* with mNeon and Sec23D with mRuby.** (A) Diagrams illustrate the result of homologous recombination-mediated insertion of mRuby into the Sec23D locus (top) and HDR-mediated insertion of mNeon into the *Sec23B* (middle) and *Sec23G* (bottom) loci. Coding exons are indicated by thick boxes and untranslated exons are indicated by thin boxes. Thin lines indicate intronic regions and thin dotted lines are intronic regions downstream of the gene. Inserted sequences are denoted by thick colored boxes (magenta, mRuby; yellow, hygromycin resistance cassette; light green, mNeon). The dashed vertical lines indicate the junction between the knock-in construct and upstream and downstream genomic sequences. Small arrows above the diagrams represent primers used for genotyping. Scale bar is 0.5 kb. (B) PCR products obtained with the indicated primer pairs using the template DNA isolated from the indicated moss line were separated on an agarose gel and stained with ethidium bromide. Molecular weight is indicated in kb. Predicted sizes for correct products are indicated in (A).

### Supplemental Tables

**Table S1. Sequence comparisons of fragments used in RNAi constructs.**

| Gene Pair | Sequence Length (bp) | Percent identity |
|---|---|---|
| Sec23B / Sec23C | 595 | 82.5 |
| Sec23D / Sec23E | 690 | 84.0 |
| Sec23F / Sec23G | 843 | 85.3 |
| Sec24A / Sec24B | 718 | 81.5 |
| Sec24C / Sec24D | 691 | 100 |
| Sec24F / Sec24G | 735 | 87.7 |

**Table S2. Mutants used in this study.**

| **Mutant** | **Background** | **Mutations** | **Translation** |
|---|---|---|---|
| *Δsec23d-5* | WT/ Sec23B-3XmRuby/Sec23G-3XmRuby | Sec23D: 11bp deletion | Stop at AA43 |
| *Δsec23d-10* | WT | Sec23D: 2 bp deletion | Stop at AA46 |
| *Δsec23d-11* | WT | Sec23D: 7 bp deletion | Stop at AA48 |
| *Δsec23d-17* | WT | Sec23D: 16 bp deletion | Stop at AA45 |
| *Δsec23d-B11* | ER-GFP | Sec23D: 7bp deletion | Stop at AA48 |
| *Δsec23d-F12* | YFP-Golgi | Sec23D: 13bp deletion | Stop at AA46 |
| *Δsec23d-S2* | F-SNAP-mCherry NLS4-4 | Sec23D: 15bp replaced with random 11bp | Stop at AA49 |
| *Δsec23d-CG7* | Sec24C-GFP | Sec23D: 29bp deletion | Stop at AA37 |
| *Δsec23e-6* | WT | Sec23E: 5 bp deletion | Stop at AA123 |
| *Δsec23afg-J4* | WT | Sec23A: 4 bp deletion<br>Sec23F: 1 bp insertion<br>Sec23G: ~3000 bp insertion | Stop at AA28<br>Stop at AA31<br>STOP*at AA22 |
| *Δsec23bg-J9* | WT | Sec23B: 19 bp deletion<br>Sec23G: 10 bp deletion | Stop at AA65<br>Stop at AA98 |
| *Δsec23acfg-K1* | *Δsec23afg-J4* | Sec23A: 4 bp deletion<br>Sec23C: 8 bp deletion<br>Sec23F: 1 bp insertion<br>Sec23G: ~3000 bp insertion | Stop at AA28<br>Stop at AA10<br>Stop at AA31<br>STOP*at AA22 |
| *Δsec23abcfg-1* | *Δsec23acfg-K1* | Sec23A: 4 bp deletion<br>Sec23B: 4 bp deletion<br>Sec23C: 8 bp deletion<br>Sec23F: 1 bp insertion<br>Sec23G: ~3000 bp insertion | Stop at AA28<br>Stop at AA53<br>Stop at AA10<br>Stop at AA31<br>STOP*at AA22 |

| | | | |
|---|---|---|---|
| *Δsec23abcfg-2* | *Δsec23bg-J9* | Sec23A: 2 bp deletion<br>Sec23B: 19 bp deletion<br>Sec23C: 1 bp deletion<br>Sec23F: 13 bp deletion<br>Sec23G: 10 bp deletion | Stop at AA50<br>Stop at AA65<br>Stop at AA42<br>Stop at AA53<br>Stop at AA98 |

*Sec23G cDNA isolated from this line has a 2296 bp deletion of the Sec23G coding sequence.

**Table S3. Gene IDs for *Sec23* and *Sec24* gene family members.**

| **Gene Name** | **Gene ID (Phytozome)** |
|---|---|
| *PpSec23A* | Pp1s77_4 |
| *PpSec23B* | Pp1s176_120 |
| *PpSec23C* | Pp1s1_345 |
| *PpSec23D* | Pp1s55_55 |
| *PpSec23E* | Pp1s181_124 |
| *PpSec23F* | Pp1s62_159 |
| *PpSec23G* | Pp1s196_14 |
| *PpSec24A* | Pp1s358_10 |
| *PpSec24B* | Pp1s570_1 |
| *PpSec24C* | Pp1s59_281 |
| *PpSec24D* | Pp1s59_293 |
| *PpSec24E* | Pp1s30_364 |
| *PpSec24F* | Pp1s29_149 |
| *PpSec24G* | Pp1s220_95 |

**Table S4. Primers used in this study.**

| Primer Name | Sequence | Purpose |
|---|---|---|
| Sec23aRNAi-F | CACCATGGATTTCACCGAGTTGG | CDS RNAi |
| Sec23aEcoRIRNAI-R | AAGGAATTCACTAATAAGCCCCACGAGTGCC | CDS RNAi |
| Sec23cBamHIRNAi-F | CACCGGATCCCCGAATGGTTTGCAGAAGTGG | CDS RNAi |
| Sec23cXbaIRNAi-R | AAGTCTAGATCCTTGGTCACATCCTTGG | CDS RNAi |
| Sec23dEcoRIRNAi-F | CACCGAATTCATGGCGGAGTTTTTGGACGC | CDS RNAi |
| Sec23dBamHIRNAi-R | AAGGGATCCGCCAGCGGATTGAAGACCAAGC | CDS RNAi |
| Sec23gXbaIRNAi-F | CACCTCTAGAGTACACGACGCCCTGGTTTTGTG | CDS RNAi |
| Sec23gRNAi-R | CTATTACACGAGTCACGGCAACG | CDS RNAi |
| Sec23d-5UTR-F | CACCAGCGTTGGCGATTGAACGTCAACG | UTR RNAi |
| Sec23d-5UTR-BamHI-R | AAGGATCCGCAACCCCTCTCGCAACTCTATGA | UTR RNAi |
| Sec23d-3UTR-BamHI-F | CACCGGATCCAGTTATGATGGAGATATACTGC | UTR RNAi |
| Sec23d-3UTR-R | CCAACTTTGAAACAAGCTGT | UTR RNAi |
| Sec23e-BamHI-5UTR-F | CACCGGATCCGCTCAGTTGAACTTTGAGGAGC | UTR RNAi |
| Sec23e-EcoRI-5UTR-R | AAGAATTCCGAGAGCAATGCTCCTAAACC | UTR RNAi |
| Sec23e-EcoRI-3UTR-F | CACCGAATTCTTGGTCACCTAAGATCCTCACC | UTR RNAi |
| Sec23e-BamHI-R | AAGGATCCTCAAATTCTGGAAGATATGGGAAAAAA | UTR RNAi |
| attB1-Sec23d-F | GGGGACAAGTTTGTACAAAAAAGCAGGCTTTATGGCGGAGTT TTTGGACG | Amplify cds |
| attB5r-Sec23d-R | GGGGACAACTTTTGTATACAAAGTTGTAGACTGCACTACTGCT AGCC | Amplify cds |
| CACC-Sec23d-F | CACCATGGCGGAGTTTTTGGACG | Amplify cds |
| Sec23d-R | TCAAGACTGCACTACTGCTAG | Amplify cds |
| CACC-Sec23e-F | CACCATGGCTGAATTTGTGGAAGC | Amplify cds |
| Sec23e-R | TCAAGATTGGACAACTGCC | Amplify cds |
| Se-Sec23dCDS-F | CGAATTAGGGTTTTCGGATTTTGG | Sequencing |
| Se-Sec23eCDS-F | GGTTTTCTGAATTTGGTTGTCC | Sequencing |
| Se-Sec23deCDS-R | TCCCCAAACCGCTGTGAGA | Sequencing |
| Sec23D-attB1-SwaI-F | GGGGACAAGTTTGTACAAAAAAGCAGGCTATTTAAATTAACTG CTCCAAGGATGTTAAG | Tagging |
| Sec23D-attB5r-R | GGGGACAACTTTTGTATACAAAGTTGTAGACTGCACTACTGCT AG | Tagging |
| Sec23D-attB3-F | GGGGACAACTTTGTATAATAAAGTTGACTGTGTTCATTTTACTT GCGG | Tagging |
| Sec23D-attB2-SwaI-R | GGGGACCACTTTGTACAAGAAAGCTGGGTAATTTAAATAGGG GCAGGAGGCT | Tagging |
| Sec23D-crispr-F | CCATGCGTGTAGAGCGCCGCGAGT | Sec23D crispr |
| Sec23D-crispr-R | AAACACTCGCGGCGCTCTACACGC | Sec23D crispr |
| Sec23E-crispr-F | CCATTACGTGCCCCCGCCCATGCC | Sec23E crispr |
| Sec23E-crispr-R | AAACGGCATGGGCGGGGGCACGTA | Sec23E crispr |
| Sec23D-tagging-up-F | TTTCTTGCGGCTCTTTTAATCAAC | Genotyping |

| | | |
|---|---|---|
| NOSter-jct-Rev | ATGCTTAACGTAATTCAACAG | Genotyping |
| QuadGateHygroForw | CTCTAGAGTCGAGGGTACG | Genotyping |
| Sec23D-tagging-down-R | CCAAAGATACATGAAGAATCGCAA | Genotyping |
| Sec23D-CRIPSR-GF | AGTGTGAACAGAAGTTTTTTTCG | Genotyping |
| Sec23D-CRISPR-GR | CCTTAGAAAATCCAAAATCCGAA | Genotyping |
| Sec23E-CRISPR-GF | TCGTTTAATTAGGCATCGTTTTG | Genotyping |
| Sec23E-CRISPR-GR | GCTCAGGTGCTAAGTAAAAC | Genotyping |
| Sec23A-crispr-F | CCATGAGTCGCGTAGCCTCAATCC | Sec23A crispr |
| Sec23A-crispr-R | AAACGGATTGAGGCTACGCGACTC | Sec23A crispr |
| Sec23B-crispr-F | CCATGGGTAGCACATTCACCGGTG | Sec23B crispr |
| Sec23B-crispr-R | AAACCACCGGTGAATGTGCTACCC | Sec23B crispr |
| Sec23C-crispr-F | CCATCTACGTGTTTCGTGGCGCCA | Sec23 crispr |
| Sec23C-crispr-R | AAACTGGCGCCACGAAACACGTAG | Sec23 crispr |
| Sec23F-crispr-F | CCATCTCTCGCCCGACCCAGCAGG | Sec23 crispr |
| Sec23F-crispr-R | AAACCCTGCTGGGTCGGGCGAGAG | Sec23 crispr |
| Sec23G-crispr-F | CCATAGGATCTCTTGACCCGGCGA | Sec23 crispr |
| Sec23G-crispr-R | AAACTCGCCGGGTCAAGAGATCCT | Sec23 crispr |
| Sec23A crispr GF | TCAGAACTTCTGAAGAGGCGAC | Genotyping |
| Sec23B crispr GF | ACTTCGGCTCCTGTCTTTTAG | Genotyping |
| Sec23A crispr GF | TTTTCAGCCATCCCAAACTG | Genotyping |
| Sec23B crispr GR | CCTTAATTCCAGATCCCACAGTG | Genotyping |
| Sec23C crispr GR | CAATAATTCCAGATCCCACAGTG | Genotyping |
| Sec23G crispr GR | AGCTCCTAACCCTTCATCA | Genotyping |
| Sec23G crispr GF | AAACCATGACGACGACGAA | Genotyping |
| Sec23F crispr GR | ACTTCCTACATCGGTCATATACAC | Genotyping |
| Sec23F crispr GF | ACCAAGTTAGTGCAGGTGATG | Genotyping |
| PPBip1-qF | GGAAGTCGGAGAAGATCGTG | qRT-PCR |
| PPBip1-qR | CCCAGCTGGTTCTTCATGTT | qRT-PCR |
| Sec23A-qF | ATGTTCTGTCGCATCCCCTG | qRT-PCR |
| Sec23A-qR | AGTCAAGTCCTCCTCGGTCA | qRT-PCR |
| Sec23B-qF | AGCACGAAGAACATACGGCA | qRT-PCR |
| Sec23B-qR | AAACGAGCTTGCGACCCATA | qRT-PCR |
| Sec23C-qF | GACCGAATCCTGCTGTTGGA | qRT-PCR |
| Sec23C-qR | GGGAACCGCTCGTGGATAAT | qRT-PCR |
| Sec23D-qF | GAGAGGTTCTTGCGGTCAGT | qRT-PCR |
| Sec23D-qR | CTTTGAGGGCTGCACCAGTA | qRT-PCR |
| Sec23E-qF | TATCCGTAGCCGATTGCGAG | qRT-PCR |
| Sec23E-qR | TGCAGGGTCCGCCAATAAAA | qRT-PCR |
| Sec23F-qF | CGAGATGTTGCTGACATGCG | qRT-PCR |

| | | |
|---|---|---|
| Sec23F-qR | TGACAAATCGAAGCCAGCCT | qRT-PCR |
| Sec23G-qF | AGCTTCCAGTGAGCACATCC | qRT-PCR |
| Sec23G-qR | ACAGGTCCCAGCACACAAAA | qRT-PCR |
| Sec23D-CRIPSR-GF_UM2397 | AGTGTGAACAGAAGTTTTTTTCG | Genotyping |
| Sec23D-CRISPR-GR_UM2398 | CCTTAGAAAATCCAAAATCCGAA | Genotyping |
| Sec23D-XmaI-R | TACACTGCGGCCAAATCTCCC | Sequencing |
| EGFP-int-F | GGCATCAAGGTGAACTTCAAGATCC | Sequencing |
| EGFP-int-F2 | CCTACGGCGTGCAGT | Sequencing |
| EGFP-int-R2 | GGATCTTGAAGTTCACCTTGATGCC | Sequencing |
| Sec23D-crispr2-F | CCATACCGCTACGGGATTGCTATG | Sec23D crispr |
| Sec23D-crispr2-R | AAACCATAGCAATCCCGTAGCGGT | Sec23D crispr |
| Sec23G-crispr-GUR | TTCGTCGTCGTCATGGTTT | Sec23G crispr |
| Sec23G-crispr-GDF | TGATGAAGGGTTAGGAGCT | Sec23G crispr |
| Sec23G crispr CACC-GF | CACCAAACCATGACGACGACGAA | Sec23G crispr |
| Sec23D crispr CACC-GF | CACCAGTGTGAACAGAAGTTTTTTTCG | Sec23D crispr |
| Sec23G-crispr-2F | CCATCCGCCTCCGACGTTTGGTGC | Genotyping |
| Sec23G-crispr-2R | AAACGCACCAAACGTCGGAGGCGG | Genotyping |
| Sec23G-HDR-GF | CACCCGTAAAATTTCGCCACG | Genotyping |
| Sec23G-HDR-GR | GAATCTCTGCGAAATTGGAAGG | Genotyping |
| Sec23G-crispr-G2R | GCAAAGTGAACGGGAATTGT | Genotyping |
| Sec23G-crispr-G3R | CGTGGCGAAATTTTACGGGTG | Genotyping |
| Sec23B-crispr-HDR-F | CCATATCATCTAGTTGTGCTGTGG | Sec23B HDR CRISPR |
| Sec23B-crispr-HDR-R | AAACCCACAGCACAACTAGATGAT | Sec23B HDR CRISPR |
| Sec23B-UpattB1-F | GGGGACAAGTTTGTACAAAAAAGCAGGCTTACACTACGAAGGAGATATGCGGC | Tagging |
| Sec23B-UpattB4-R | GGGGACAACTTTGTATAGAAAAGTTGGGTGGTTTTGCACAGCCAGCCTTTTCAA | Tagging |
| Sec23B-DownattB3-F | GGGGACAACTTTGTATAATAAAGTTGATTTGTGCTGTGGTGGACTCTACTTG | Tagging |
| Sec23B-DownattB2-R | GGGGACCACTTTGTACAAGAAAGCTGGGTAGTTCCAATCTCCTTTCATAATTCACCCAA | Tagging |
| Sec23G-crispr-HDR-F | CCATCCCGAGACATATACACGTTT | Sec23G HDR CRISPR |
| Sec23G-crispr-HDR-R | AAACAAACGTGTATATGTCTCGGG | Sec23G HDR CRISPR |
| Sec23G-UpattB1-F | GGGGACAAGTTTGTACAAAAAAGCAGGCTTAGTAAATGGTGGCACCTACATGAGT | Tagging |

| | | |
|---|---|---|
| Sec23G-UpattB4-R | GGGGACAACTTTGTATAGAAAAGTTGGGTGAACACTAAGCTTAAGGCGTCTCATCC | Tagging |
| Sec23G-DownattB3-F | GGGGACAACTTTGTATAATAAAGTTGATTACACGTTTTGGTAAATATAGTCCACATATTGGTG | Tagging |
| Sec23G-DownattB2-R | GGGGACCACTTTGTACAAGAAAGCTGGGTATGGTAGGTGGTAGATGGCAAGTTGTG | Tagging |
| Sec23B-HDR-GF | GGAAATTGAGCGGCTTGGA | Genotyping |
| mNeon-Int-F | GGCTATGGTAGATGGCAGT | Genotyping |
| Sec23G-HDR-GF | TGTTCAGGTGATCGGATGGTTC | Genotyping |
| Sec23G-HDR-GR | TCCTGAAGTTGTGTTGCAC | Genotyping |
| Sec24A qF1 | GTAGTGGAACAGGAGACCGC | qRT-PCR |
| Sec24A qR1 | GGGTTGTTCCCCTTGCCTAA | qRT-PCR |
| Sec24B qF1 | ACTGCTGAGGTTGCTGTACC | qRT-PCR |
| Sec24B qR1 | AAGAACCAAAGGAGCAGGCA | qRT-PCR |
| Sec24C qF1 | CTTTTTGGCGTGTGTTCGGT | qRT-PCR |
| Sec24C qR1 | ATGCACAACCGCAGGTATGA | qRT-PCR |
| Sec24D qF1 | CTTCCAACTCAGGCAGACCA | qRT-PCR |
| Sec24D qR1 | TGTTGTTGGTACGGCAAGGA | qRT-PCR |
| Sec24E qF1 | CACCTGGGGAGGAAGTTACAG | qRT-PCR |
| Sec24E qR1 | CACGAGAAAGCTCTGGCCTT | qRT-PCR |
| Sec24F qF1 | GACACTAGACAAGCCAGCGT | qRT-PCR |
| Sec24F qR1 | TTTTGAGGTGGTGGGAACGG | qRT-PCR |
| Sec24G qF1 | GCTGAAGCAGGAGAGGAACA | qRT-PCR |
| Sec24G qR1 | CAAGAACGAAACGAACCCCAC | qRT-PCR |
